# Crosstalk between and developmental dynamics of *C. elegans* Argonaute proteins

**DOI:** 10.64898/2025.12.17.694999

**Authors:** Ann-Sophie Seistrup, Emily Nischwitz, Falk Butter, René F. Ketting

## Abstract

Gene regulation via Argonaute-bound small RNAs is a broadly conserved mechanism, present in all domains of life. The nematode *Caenorhabditis elegans* expresses several worm-specific Argonautes (WAGOs), which interact with the small RNAs known as 22G-RNAs. These WAGOs have roles in gene regulation, transposon-defence as well as in viral control and experimentally induced RNA interference. Despite many studies, direct relationships between WAGO targeting, as defined by 22G RNA sequences, and mRNA abundance are not clear. Also, the effects of developmental stage and WAGO-interconnectivity have been under-studied thus far. We studied these aspects for two germline-expressed WAGO proteins, WAGO-1 and WAGO-3. We show that WAGO-1 mostly affects 22G-RNA expression in gravid adult worms, while WAGO-3 predominantly affects 22G-RNA expression in embryos. Furthermore, we detect a link between WAGO-3 and the maternal 26G-RNA pathway governed by the Argonaute protein ERGO-1, and between WAGO-1 and the paternal 26G-RNA pathway governed by the Argonautes ALG-3/4. We also demonstrate that, globally speaking, loss of WAGO-1 or WAGO-3 does not result in upregulation of their target mRNAs, as defined by 22G-RNA complementarity. Finally, metagene analysis of 22G-RNA profiles suggests loss of one WAGO protein leads to shifts in WAGO 22G-RNA binding. Overall, we conclude that WAGO-1 and WAGO-3 are developmentally dynamic, are embedded in distinct regulatory networks, and that potential silencing of individual mRNAs by these WAGO proteins is hard to assess by simple loss-of-function studies.

**Summary:** RNA interference describes silencing of genes via small RNAs coupled to an Argonaute protein. *C. elegans* expresses a class of secondary small RNAs known as 22G-RNAs which are coupled to Argonautes known as WAGOs (worm-specific Argonautes). We have sequenced 22G-RNA and mRNA populations from deletion mutants of two such WAGOs, WAGO-1 and WAGO-3, to investigate Argonaute crosstalk and the relationship between 22G-RNA and mRNA levels. We found that WAGO-1 and WAGO-3 engage different, complementary pathways, and that target silencing cannot be deduced simply from 22G-RNA complementarity.

## Introduction

RNA interference (RNAi) describes an important mechanism for gene regulation and transposon silencing, in which a small RNA (roughly 20-30 nucleotides (nt) in length) interacts with a protein of the Argonaute family. This complex binds a target mRNA via sequence complementarity to the small RNA and can exert a variety of functions, generally leading to repression (Ha & Kim, 2014; Ketting & Cochella, 2021). Although first discovered in the late 1990s (Fire et al., 1998), much is still unknown about the exact mechanisms that guide this repression.

Several different types of small RNAs exist and they all bind distinct Argonaute proteins (Ketting & Cochella, 2021). Argonaute classification is based on sequence homology, yet all classes share the same general structure. This consists of four folded domains, named N-, PAZ-, MID-, and PIWI-domain (Wu et al., 2020). While the PAZ and MID domain aid in anchoring the small RNA, the PIWI domain adopts an RNaseH-like fold that can be catalytically active and can cleave target mRNAs. However, many Argonaute proteins have lost catalytic activity and affect their target transcripts in other ways (Czech & Hannon, 2016; Wu et al., 2020). The N-terminal region plays a role in unwinding and loading of the small RNA (Wu et al., 2020). Moreover, many Argonautes, including worm-specific Argonautes (WAGOs), contain an additional unstructured N-terminal tail, which may play a role in phase separation and subcellular localization (Uversky et al., 2015).

WAGO proteins have undergone large genetic expansion in nematodes and bind a particular class of small RNAs known as 22G-RNAs (Hoogstrate et al., 2014). 22G-RNAs are so-called secondary small RNAs, as they are induced by the activity of Argonaute proteins, so-called primary Argonautes. Because 22G-RNAs are formed by RNA directed RNA polymerases (RdRPs), with no further 5’-end processing required, they carry a triphosphate at the 5’-end, which distinguishes them from the other classes of small RNAs (Ketting & Cochella, 2021). In order to enrich for 22G-RNAs in sequencing studies, this triphosphate must first be cleaved via treatment with either Tobacco Acid pyrophosphatase (TAP) or RNA 5’ pyrophosphohydrolase (RppH) (Almeida, de Jesus Domingues, Lukas, et al., 2019).

*C. elegans* encodes 26 different Argonaute genes for which 19 have proven expression, with 11 of these belonging to the WAGO family (Seroussi et al., 2023). Primary Argonaute proteins include PRG-1, the *C. elegans* Piwi protein that binds 21U-RNAs, and ERGO-1, ALG-3 and ALG-4. The latter three define the endo-siRNA, or 26G-RNA pathway. The 26G-RNA Argonautes show differential expression throughout the development of the hermaphroditic germline, with ERGO-1 being expressed during oogenesis as well as in embryos and the mutually redundant Argonautes ALG-3 and ALG-4 being only expressed during spermatogenesis (Almeida, de Jesus Domingues, & Ketting, 2019; Conine et al., 2010; Han et al., 2009). A fifth primary Argonaute protein is RDE-1, which responds to environmental double stranded RNA and is essential for RNA interference (RNAi)(Tabara et al., 1999). The full extent of how much the Argonaute pathways interconnect is not yet fully understood (Liu et al., 2023; Seroussi et al., 2023).

Several WAGOs have been shown to play a role in RNAi and its inheritance(Yigit et al., 2006). Recently, we showed that WAGO-3 is important for inheritance both via the maternal and paternal gametes (Schreier et al., 2022; Schreier et al., 2025), and also WAGO-1 has been shown to affect RNAi (Woodhouse et al., 2025). WAGO-1 resides in the same subcellular location as WAGO-3; the P-granule, a perinuclear focus in the germline which is formed via phase-separation (Gu et al., 2009; Schreier et al., 2022). WAGO-1 and −3 share some of their targets as based on RNA-immunoprecipitation sequencing (RIP-Seq) and they both target transposons and pseudogenes in addition to protein coding genes (Gu et al., 2009; Liu et al., 2023; Seroussi et al., 2023; Vastenhouw et al., 2003). Furthermore, WAGO-1 and −3 share some similarities in their N-terminal intrinsically disorder regions (IDRs) and can both be processed by the endonuclease DPF-3 (Gudipati et al., 2021). It is currently unclear which primary Argonautes are responsible for loading WAGO-1 and WAGO-3 with 22G RNAs, and the exact mechanism of WAGO-1 and WAGO-3 enact gene silencing are also poorly understood, although comparison of protein and mRNA levels of a WAGO-1 target suggests that WAGO-1 functions in mRNA turnover (Aoki et al., 2021). We have recently demonstrated that loss of the germline Argonaute, WAGO-4, is accompanied by differential changes to 22G-RNAs and mRNAs at different life stages, and that, globally, these changes do not coincide, i.e. loss of 22G-RNAs does not generally come with a gain of mRNAs that are targeted by the affected 22G-RNAs (Seistrup et al., 2025).

Here, we have studied the closely related Argonautes, WAGO-1 and WAGO-3 (also known as PPW-2), and find that loss of these comes with mostly unrelated changes to 22G-RNA and mRNA landscapes; we detected only 18 genes, that, based on these changes, may be direct targets of WAGO-3, and just one that may be a direct target of WAGO-1. Our data also show that developmental stage is a key variable that has to be taken into account when studying WAGO protein function. Furthermore, we show that WAGO-1 and WAGO-3 affect the ALG-3/4 and ERGO-1 pathways respectively. Finally, shifts in 22G RNA metagene profiles suggest shifts in 22G RNA-WAGO-interactions upon loss of WAGO-1 or WAGO-3.

## Results

### WAGO-3 binds similar 22G-RNAs in L4 larvae and gravid adult worms

We have previously demonstrated that targets of WAGO-3 differ slightly between sexes, with the interesting observation that 22G-RNAs against the transposon Tc3 were pulled down along with GFP::3xFLAG::WAGO-3 in hermaphrodites but not in males (Schreier et al., 2022). We wanted to test whether this observation, and general WAGO-3 targeting, was specific to sex or rather to spermatogenesis. To this end, we did RNA immunoprecipitation sequencing (RIP-Seq) of GFP::3xFLAG::WAGO-3 L4 hermaphroditic larvae, which are undergoing spermatogenesis (Supplementary Figure 1). With our cut-offs (non-zero value in input, two-fold or higher enrichment in IP, and RPKM in IP 4 or higher for non-transposon genes in two or more replicates) we identified 1,475 targets in the L4. This is significantly more than what we observed before (Schreier et al., 2022) in gravid adults and males (569 and 597 respectively). We note, however, that the L4 experiment was done at a later timepoint, and stronger enrichment of 22G-RNAs in the more recent experiment (Supplementary Figure 1a) may allow the detection of more 22G-RNA targets. Hence, we would not conclude from this data that in L4 stage WAGO-3 has more targets.

Gravid adult, male, and L4 targets overlapped significantly and Tc3 was a target of WAGO-3 in L4 hermaphrodites (Figure 1b and Supplementary Figure 1c), suggesting that the loss of Tc3 targeting is a male-specific feature, and not a spermatogenesis-specific feature. The overall distribution of target types did not change between life stages and sexes, with all three cases showing transposons and pseudogenes as over-represented targets in WAGO-3 (Figure 1c).

**Figure 1:**
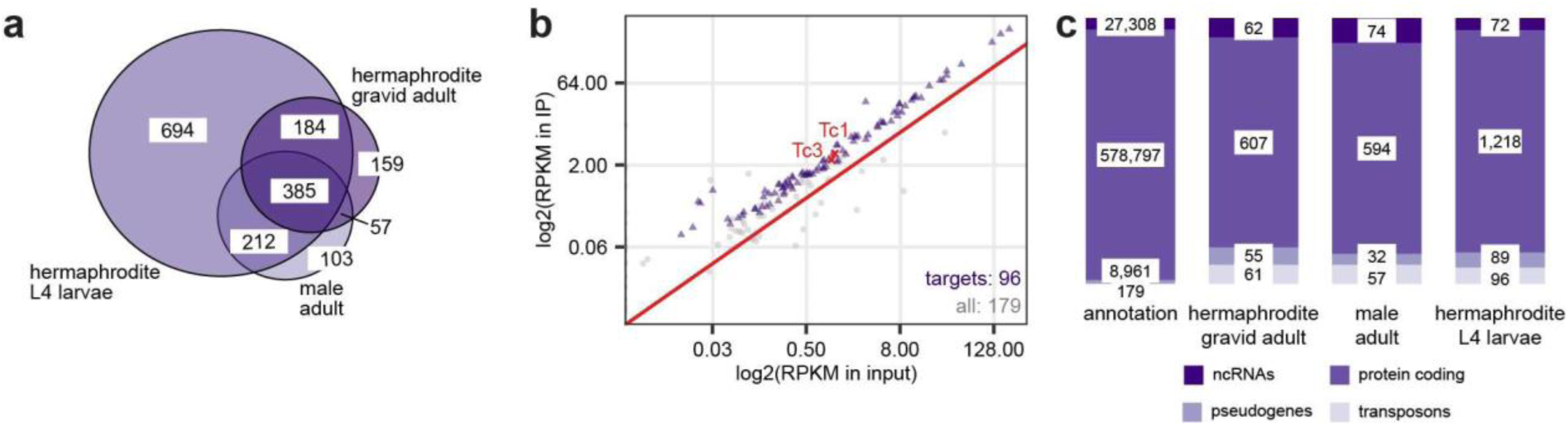
RNA-immunoprecipitation sequencing of GFP::3xFLAG::WAGO-3(*xf119*). **a)** Venn diagram showing the overlaps of WAGO-3 targets found in L4 larvae and gravid adult hermaphrodites as well as adult males. **b)** Scatterplot showing reads mapping to transposons in input and IP samples from hermaphroditic L4 larvae. Each point represents one transposon with one or multiple insertions and typically several 22G-RNAs mapping to it. **c)** Bar plot showing the distribution of gene types in the annotation and in the WAGO-3 targets in L4 hermaphroditic larvae, gravid adult hermaphroditic worms and adult male worms.

### Loss of WAGO-3 impacts embryos more than other life stages

To further probe WAGO-3 function in relation to life stage, we sequenced small RNAs from three different life stages (embryos, L4 larvae, and gravid adult worms) of hermaphroditic *wago-3(pk1763)* deletion mutants (henceforth *wago-3*). We also generated poly(A)-selected libraries from the very same samples in order to investigate mRNA changes caused by loss of WAGO-3.

First, we confirmed that WAGO-3 was not expressed in the *wago-3* samples (Supplementary Figure 2a) and noted no overall loss or gain in the relative expression of 22G-RNAs (Supplementary Figure 2b+c). 22G-RNA levels were lower in embryo libraries relative to the other life stages, but not differentially between *wago-3* and wild type (Supplementary Figure 2b+c). Nonetheless, we did note that *wago-3* embryos expressed relatively more miRNAs than wild type embryos, hinting at a global embryonic small RNA mis-regulation caused by loss of a single WAGO (Supplementary Figure 2c).

Roughly 400-1,000 genes had significant changes in 22G-RNAs between wild-type and *wago-3* animals in either of the three life stages assessed, with the fewest changes happening in the embryos (Figure 2a). At the mRNA level, L4 larvae and gravid worms responded only mildly to *wago-3* deletion, with only about 200 genes displaying significant differences in either case (Figure 2b). Much more striking was the effect on mRNAs in embryos, where more than 6,000 genes had reduced mRNA levels and almost 2,000 genes had elevated mRNA levels (Figure 2b). This reveals a significant impact of WAGO-3 on the transcriptome of embryos. As our L4 and gravid adult samples did not show these effects, they apparently balance out as the animals develops further. On both the 22G-RNA and mRNA levels, many of the significant changes were life stage specific (Supplementary Figure 3a-b).

**Figure 2:**
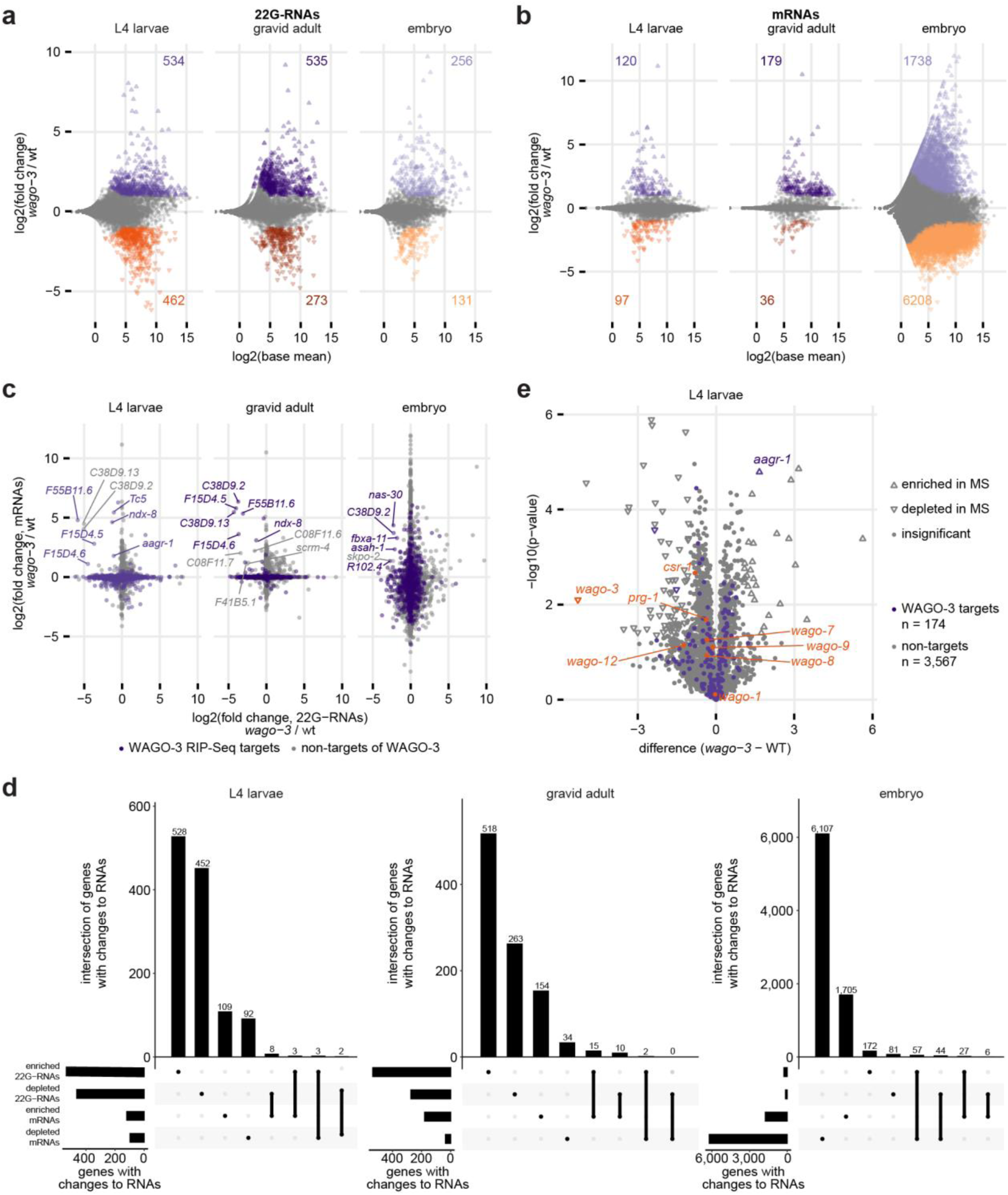
Sequencing of RNAs from *wago-3* mutant animals. **a)** Scatterplot showing changes to 22G-RNAs in *wago-3* as compared to wild type in three different life stages, as indicated. Colours represent genes with significantly depleted/enriched 22G-RNA levels; numbers indicated in each plot. Each dot represents one gene, which typically has several 22G-RNAs mapping to it. **b)** Scatterplot showing changes to mRNA levels in *wago-3* as compared to wild type in three different life stages, as indicated. Colours represent genes with significantly depleted/enriched 22G-RNA levels; numbers indicated in each plot. **c)** Scatterplot showing the comparison between 22G-RNA levels and mRNA levels for all genes detected. Genes that had significantly depleted 22G-RNA levels and significantly enriched mRNA levels are indicated. Colours represent whether each gene was defined as a target of WAGO-3 via RIP-Seq. For L4 larvae, results from RIP-Seq of L4 larvae (Figure 1) was used as the comparison. For gravid adult worms and embryos, results from RIP-Seq of gravid adult worms was used (Schreier et al., 2022). **d)** UpSet plots showing the overlap between genes with significantly enriched/depleted mRNA levels and 22G-RNA levels in each of the three life stages. **e)** Scatterplot showing changes to peptide levels in *wago-3* L4 larvae compared to wild type as determined by mass spectrometry. Purple/grey indicate whether each gene was determined to be a target of WAGO-3 in L4 larvae via RIP-Seq. Argonaute proteins are shown in orange. Any Argonaute not shown in the plot was not detected.

Genes whose mRNA expression or 22G-RNA levels were affected by loss of WAGO-3, did not correspond to previously annotated targets of WAGO-3, as defined via RIP-Seq (Supplementary Figure 3c-f). Looking at the cumulative distribution function of WAGO-3 versus non-WAGO-3 targets, we saw no general up- or downregulation of 22G-RNAs or mRNAs, as evidenced by the absence of a horizontal shift of the curve, but we did see general regulation in both directions, as demonstrated by the change in the slope of the curve (Supplementary Figure 3d+f). Moreover, there were only very few cases in which a loss of 22G-RNA was coupled to a gain in mRNA levels (Figure 2c-d), suggesting that most of the dysregulation was indirect. Nonetheless, we did find 8 genes in L4 larvae, 10 genes in gravid adult, and 6 genes in the embryos where 22G-RNA levels were significantly depleted and mRNA levels were significantly enriched, suggesting these may be directly regulated by WAGO-3 (Figure 2c). There were no unifying characteristics of these genes that we could determine. Notably, five of these genes overlapped between L4 larvae and gravid adult animals (*ndx-8*, F15D4.5, F15D4.6, C38D9.2, and C38D9.13), one of which (C38D9.13) also showed coupled 22G-RNA-mRNA regulation in the embryos.

The lack of 22G-mRNA correlation made us wonder whether the silencing effect of WAGO-3-associated 22G-RNAs might affect translation instead of RNA stability. To address this, we performed mass spectrometry of *wago-3* and wild type animals (L4 stage) to probe for changes to protein levels of known WAGO-3 targets. We were able to detect 174 of the 1,475 genes that we defined as targets of WAGO-3 via RIP-Seq in L4 larvae (12%), but these did not show consistent changes in any specific direction (Figure 2e). The one target, as based on RIP-Seq, that had increased peptide levels in the *wago-*3 mutant was *aagr-1*, which we interestingly also found to have increased mRNA levels and decreased 22G-RNA levels in *wago-3*, suggesting that this is a true, direct target of WAGO-3 in L4 larvae.

Overall, while loss of WAGO-3 triggered similar effects at the 22G-RNA level at all life stages, it triggered a much stronger effect in the embryos, revealing a specific impact of WAGO-3 on embryonic gene expression. However, the lack of a clear anti-correlation between mRNA and 22G-RNAs shows that much of this impact likely is indirect, and that reliable target-identification from *wago-3* mutant data is not possible.

### Loss of WAGO-3 causes overproduction of WAGO-6/8/12-like 22G-RNAs

The fact that loss of WAGO-3 does not cause depletion of 22G-RNAs against WAGO-3 targets, even though 22G-RNAs are expected to require WAGO-binding to be stabilized, suggests that other Argonautes may compensate the loss of WAGO-3-bound 22G-RNA sequences. While the RdRPs that synthetize the 22G-RNAs can presumably produce 22G-RNAs from any part of the target RNA, RIP-Seq experiments have revealed that different WAGOs show different metagene profiles of the 22G-RNAs with which they associate (Almeida, de Jesus Domingues, & Ketting, 2019; Charlesworth et al., 2021; Chen & Phillips, 2025; Conine et al., 2010; Jelenic et al., 2025; Schreier et al., 2022; Seroussi et al., 2023)(Figure 3a). Even though the biological relevance and origin of these profiles are currently unclear, they can be used to assess by which WAGO families specific 22G-RNA pools may be bound. The WAGO-3 profile shows a gradual increase of coverage towards the 3’ end of the genes. We saw no difference in these profiles for annotated ‘WAGO-3-type’ 22G-RNAs (as defined by RIP-seq) between *wago-3* and wild type animals (Figure 2b), suggesting that the compensating WAGO(s) share the same mapping signature as WAGO-3 itself, such as WAGO-6 (also known as SAGO-2), WAGO-7 (also known as PPW-1), and WAGO-9 (also known as HRDE-1)). Likewise, analysis of the 22G-RNAs that were depleted from *wago-3* animals, irrespective of whether they represent annotated WAGO-3-targets or not, revealed the same metagene profile as that for annotated WAGO-3-bound 22G-RNAs (Figure 3c). This suggests that these same WAGOs (WAGO-6/7/9) may be losing some of their normally bound 22G-RNAs in absence of WAGO-3. To probe this further, we analysed the known WAGO-6, WAGO-7 and WAGO-9-bound RNAs. Contrary to our expectation, we observed a general elevation of WAGO-6-bound 22G-RNAs in L4 larvae and gravid adult animals (Figure 3d), although there was no consistent loss of mRNA from WAGO-6 targets (Supplementary Figure 4a). WAGO-7 class 22G-RNAs remained largely unchanged (Supplementary Figure 4b) whereas WAGO-9 class 22G-RNAs were depleted, most notably in L4 larvae (Supplementary Figure 4c). Finally, *wago-6* mRNA was not increased in L4 or gravid adult *wago-3* mutants, but it was increased in embryos (Figure 3g). WAGO-7 mRNA was unchanged in all life stages and WAGO-9 mRNA was decreased in embryos (Figure 3g). Our interpretation of these results is that in absence of WAGO-3, WAGO-6 levels incxrease, together with its bound 22G-RNAs, and that at the same time WAGO-9 decreases. This regulation happens in embryos but the changes to 22G-RNA homeostasis remains after the proteins themselves have returned to wild type levels.

**Figure 3:**
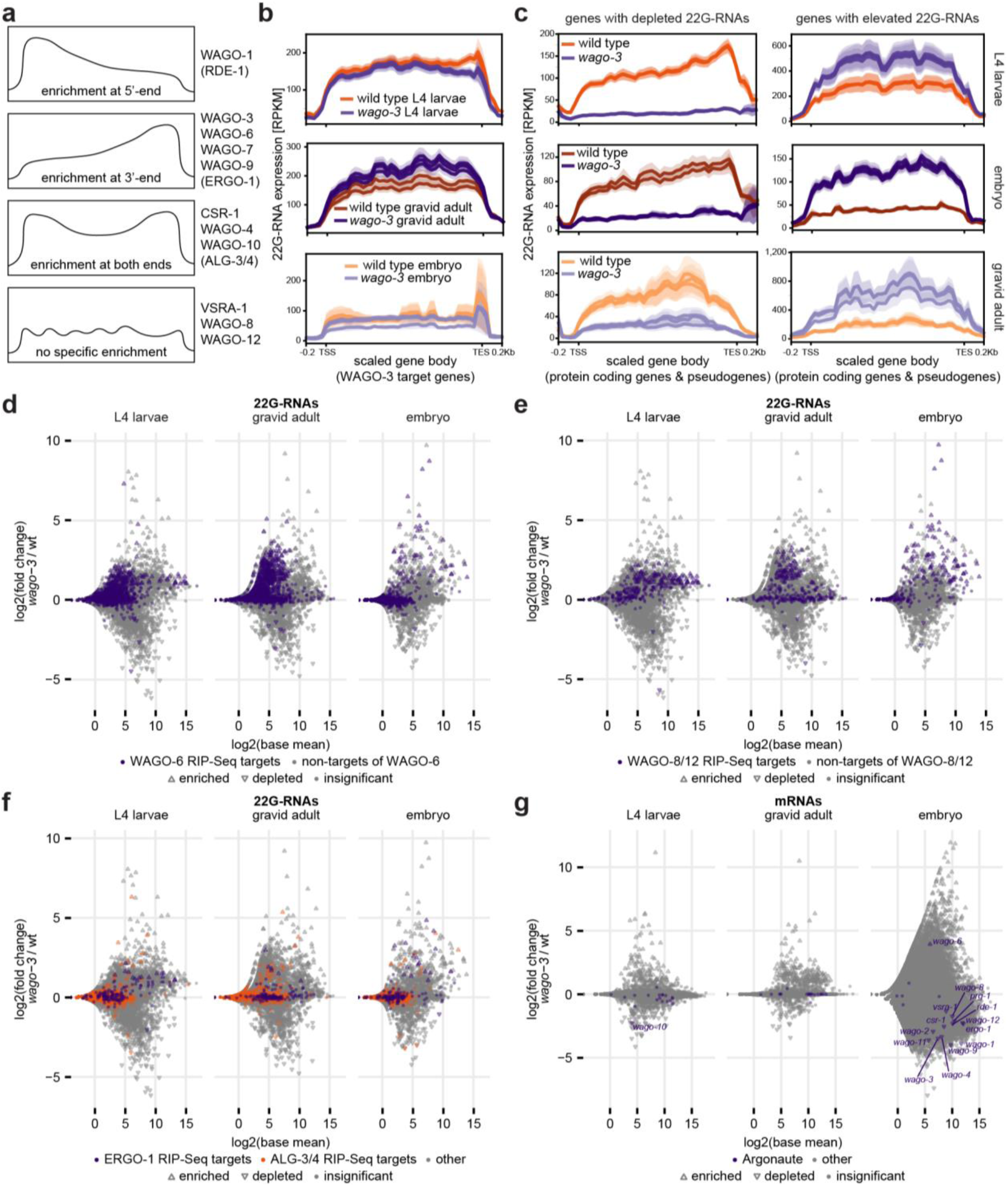
Trends of 22G-RNAs undergoing changes in *wago-3* mutant animals. **a)** Schematic of metagene profiles of different 22G-RNA binding as determined by RIP-Seq studies. Metagene profiles from primary exo-/endo-siRNA binding Argonautes are indicated in parentheses. Note that these represent general trends and that the actual data may show slight variations. **b)** Average RPKMs of 22G-RNAs along the gene body of their corresponding genes (protein coding or pseudogenes) for genes found to be targets of WAGO-3 in *wago-3* and wild type samples. For L4 larvae, results from RIP-Seq of L4 larvae (Figure 1) was used as the comparison. For gravid adult worms, results from RIP-Seq of gravid adult worms was used (Schreier et al., 2022). Embryo samples were compared to targets defined in either of these two life stages. **c)** Average RPKMs of 22G-RNAs along the gene body of their corresponding genes (protein coding or pseudogenes) for genes with elevated/depleted 22G-RNA levels in *wago-3* embryo, L4 larvae, and gravid adult animals. **d)** Scatterplot showing changes to 22G-RNAs in *wago-3* as compared to wild type in three different life stages, marked for WAGO-6 RIP-Seq targets as defined by Seroussi et al. (2023). Each dot represents one gene, which typically has several 22G-RNAs mapping to it. **e)** Scatterplot showing changes to 22G-RNAs in *wago-3* as compared to wild type in three different life stages, marked for RIP-Seq targets of WAGO-8 and/or WAGO-12 as defined by Seroussi et al. (2023). Each dot represents one gene, which typically has several 22G-RNAs mapping to it. **f)** Scatterplot showing changes to 22G-RNAs in *wago-3* as compared to wild type in three different life stages, marked for RIP-Seq targets of ERGO-1 or ALG-3/4 as defined by Seroussi et al. (2023). Overlapping genes and targets of neither are shown in grey. Each dot represents one gene, which typically has several 22G-RNAs mapping to it. **g)** Scatterplot showing changes to mRNA levels in *wago-3* as compared to wild type in three different life stages. Argonaute genes are indicated in purple; any with statistically significant changes to mRNA levels are labelled.

Next, we analysed the profiles of 22G-RNAs that were elevated in *wago-3* mutant libraries as a group. Despite the fact that WAGO-6 22G-RNAs were upregulated (see above), and that these have the same metagene profile as the WAGO-3-bound 22G RNAs, this group as a whole did show a shift in profile. These profiles were more reminiscent of WAGO-8 or WAGO-12 (also known as NRDE-3) (Figure 3c), characterised by a more or less even distribution across the gene bodies. WAGO-8 and WAGO-12 share 63-76% of their targets, and annotation of the known WAGO-8 and WAGO12 22Gs RNAs in our 22G RNA plots revealed a general increase of these22G RNAs in *wago-*3 mutants in all three life stages (Figure 3e). There was no corresponding increase of mRNAs from the *wago-8* and *wago-12* genes themselves in any life stage (Figure 3g), and peptide levels of the two proteins were also unchanged in the mass spectrometry data from L4 larvae (Figure 2e).

Finally, we noted an interesting effect on ERGO-1 targets, the maternal 26G-RNA pathway. We noted that the more highly expressed 22G-RNAs downstream of ERGO-1 increased in L4 larvae (Figure 3f), while this effect was less clear in embryos and gravid adults. As ERGO-1 is mostly expressed maternally, and L4 animals execute spermatogenesis, not oogenesis, this signal could represent erroneous ERGO-1 activation in *wago-3* mutants. In contrast, we did not detect a consistent effect on 22G-RNAs downstream of the 26G-RNA pathway that is normally expressed during spermatogenesis, i.e. the ALG-3/4 pathway (Figure 3f).

Like we observed before, these effects on 22G-RNAs didn’t translate to corresponding changes on mRNA level: neither WAGO-8/12 targets nor targets of either 26G-RNA pathway were consistently downregulated in L4 larvae or gravid adult worms (Supplementary Figure 4d). In embryos, where mRNA levels of several thousand genes were depleted, many targets of the ERGO-1 and WAGO-8/12 pathways did display reduced mRNA levels (Supplementary Figure 4d), potentially consistent with the increase of 22G-RNAs downregulating their expression. However, we note that the overall trend was towards downregulation, irrespective of WAGO-targeting (Figure 2b), suggesting that many of these changes may not reflect WAGO-targeting but rather the general trend in gene expression.

We conclude that loss of WAGO-3 leads to shifts in WAGO activities, as evidenced by changes in 22G RNA patterns. Clearly, the relationships between the different WAGO proteins are complex and cannot be resolved at this point. Yet, our results suggest connections between WAGO-3, WAGO-6, WAGO-8/12 and ERGO-1.

### GO term analysis of changing mRNAs supports role for WAGO-3 in embryogenesis

GO-term analysis of the genes with lowered mRNA levels in the embryos, revealed terms that were related to reproduction, mitosis, and embryo development (Supplementary Figure 5a), which supports the role of WAGO-3 in embryonic development. The analysis also indicated general mis-regulation of cell-cycle processes and development of organs, cytoskeleton, and nervous system (Supplementary Figure 5a). The upregulated genes were more related to defence responses and innate immunity (Supplementary Figure 5b). Even if many of these effects are indirect (see above) this analysis does show that absence of WAGO-3 leads to a lowered expression of developmental genes and a higher expression of defence-related genes. Given WAGO-3s role in transposon control, we hypothesise that these effects stem from an uncontrolled expression of ‘non-self’ genes, which in turn may induce other defence systems at the expense of the regular embryonic gene expression program.

### Loss of WAGO-1 causes large changes to mRNA levels in adults, but has no effect in embryos

WAGO-3 is found in the same perinuclear foci as WAGO-1, but the two Argonautes only share 10-25% of their targets (Seroussi et al., 2023), their metagene profiles are different (Figure 3a) and they have been proposed to have different roles (Schreier et al., 2025). To further probe these differing roles, we also included a *wago-1(ok1074)* deletion mutant, henceforth *wago-1*, in our studies. We first checked *wago-1* coverage in our mRNA-Seq data. Surprisingly, we found reads from the proposedly deleted region in our mRNA-Seq libraries (Supplementary Figure 6a). Using PCR amplification covering different regions of *wago-*1, we determined that these likely stem from insertion of the deleted region into a different genomic location (Supplementary Figure 6b-c). While we did not further assess this translocation event, our other analyses do show strong effects, indicating that *wago-1(ok1074)* is a loss of function allele, but we cannot exclude residual activity.

As for the *wago-3* strain, we did not note any overall changes to the proportional small RNA landscape of our *wago-1* mutant samples as compared to wild type (Supplementary Figure 6d-e). Only around 200 genes had significantly changed 22G-RNA levels in both *wago-1* L4 larvae and gravid adult worms (Figure 4a) and 22G-RNA levels were practically unchanged in *wago-1* embryos (Figure 4a-b). At the mRNA level, embryos were again almost unaffected, and 439 genes were upregulated in *wago-1* L4 larvae. In contrast, in *wago-1* gravid adult worms we detected almost 1,500 differentially expressed genes, of which the vast majority, 1,396 genes, had increased mRNA levels (Figure 4b). Much like in *wago-3* mutant animals, RNA changes in *wago-1* mutants were mostly life stage-specific (Supplementary Figure 7a-b), and the changes in 22G-RNA and mRNA levels appeared to be mostly non-correlative (Figure 4c-d). Only one gene, *C06A1.3*, had reduced 22G-RNA levels and elevated mRNA levels (Figure 4c), consistent with it being a direct WAGO-1 target. Notably, annotated targets of WAGO-1 as a group did not respond to the *wago-1* deletion, neither at the 22G RNA level nor the mRNA level (Supplementary Figure 7c-f). The only exception to this was that 22G-RNAs against cognate WAGO-1 targets were in fact elevated in *wago-1* L4 larvae (Supplementary Figure 7c-f). We conclude that, at least this *wago-1* allele, we cannot call direct WAGO-1-mRNA interactions.

**Figure 4:**
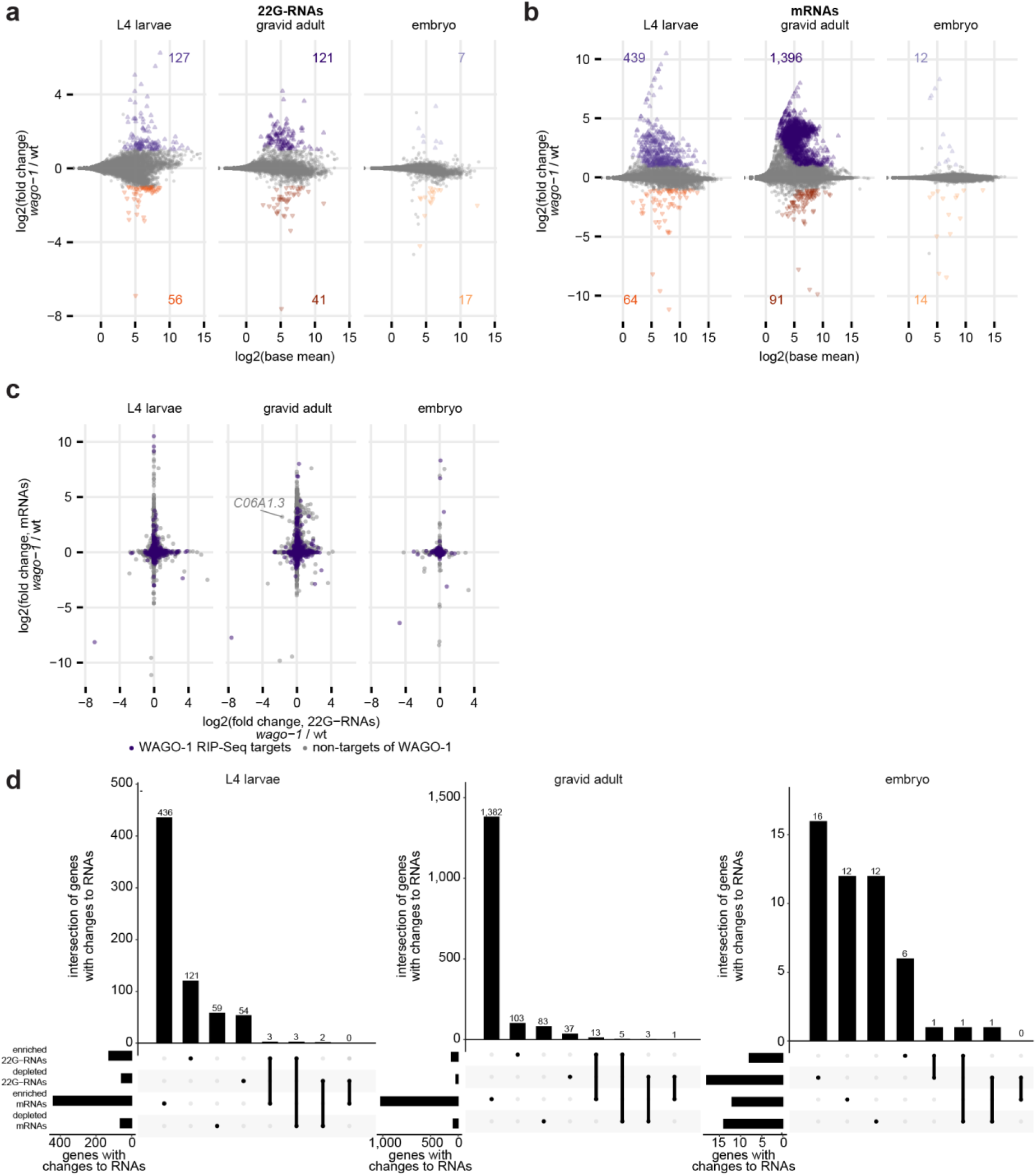
Sequencing of RNAs from *wago-1* mutant animals. **a)** Scatterplot showing changes to 22G-RNAs in *wago-1* as compared to wild type in three different life stages, as indicated. Colours represent genes with significantly depleted/enriched 22G-RNA levels; numbers indicated in each plot. Each dot represents one gene, which typically has several 22G-RNAs mapping to it. **b)** Scatterplot showing changes to mRNA levels in *wago-1* as compared to wild type in three different life stages, as indicated. Colours represent genes with significantly depleted/enriched 22G-RNA levels; numbers indicated in each plot. **c)** Scatterplot showing the comparison between 22G-RNA levels and mRNA levels for all genes detected. Genes that had significantly depleted 22G-RNA levels and significantly enriched mRNA levels are indicated. Colours represent whether each gene was defined as a target of WAGO-1 via RIP-Seq (Seroussi et al., 2023). **d)** UpSet plots showing the overlap between genes with significantly enriched/depleted mRNA levels and 22G-RNA levels in each of the three life stages.

### WAGO-1 is connected to the paternal 26G-RNA pathway

The 22G-RNAs that were lost in *wago-1* mutant animals did not reveal the WAGO-1-type metagene profile, which is enrichment at the 5’ end of the gene (Figure 5a). Rather, the mapping signature of these 22G-RNAs resembled that of WAGO-8, WAGO-12, or VSRA-1, *i.e.* spread out over the gene body. In the L4 larvae, this was presumably due to disruptions to VSRA-1, as 22G-RNAs against VSRA-1 target genes were depleted from our *wago-1* samples in L4 larvae, whereas 22G-RNAs against WAGO-8/12 target genes were unchanged or even slightly enriched (Figure 5b). As we observed before, these changes in 22G-RNAs did not translate to corresponding changes at the mRNA level (Supplementary Figure 8a).

**Figure 5:**
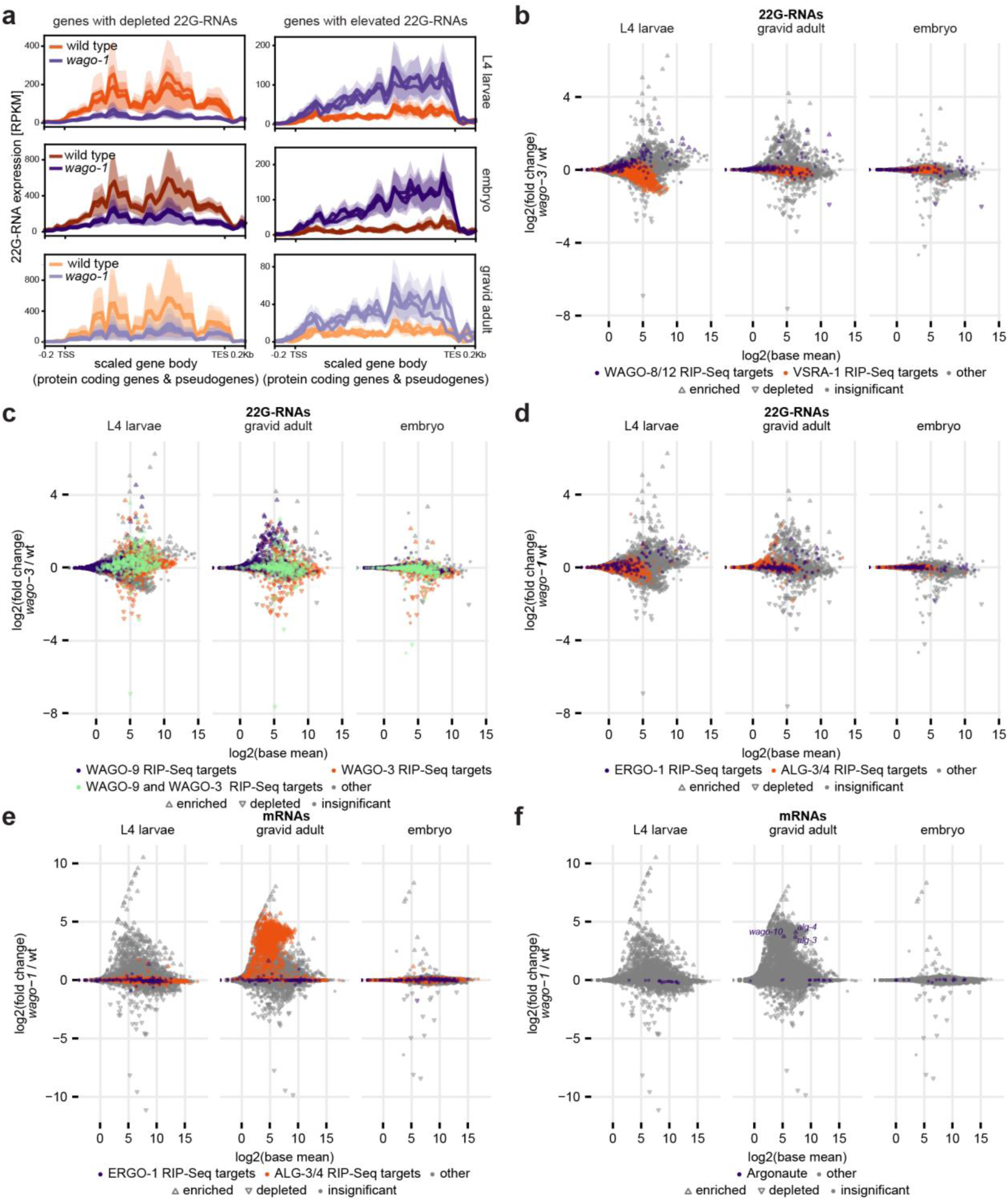
Trends of 22G-RNAs and mRNAs undergoing changes in *wago-1* mutant animals. **a)** Average RPKMs of 22G-RNAs along the gene body of their corresponding genes (protein coding or pseudogenes) for genes with elevated/depleted 22G-RNA levels in *wago-1* embryo, L4 larvae, and gravid adult animals. **b)** Scatterplot showing changes to 22G-RNAs in *wago-1* as compared to wild type in three different life stages, marked for RIP-Seq targets of WAGO-8/12 or VSRA-1 as defined by Seroussi et al. (2023). Overlapping genes and targets of neither are shown in grey. Each dot represents one gene, which typically has several 22G-RNAs mapping to it. **c)** Scatterplot showing changes to 22G-RNAs in *wago-1* as compared to wild type in three different life stages, marked for RIP-Seq targets of WAGO-9 (Seroussi et al., 2023) and/or WAGO-3 (Schreier et al., 2022 and this study; combined). Each dot represents one gene, which typically has several 22G-RNAs mapping to it. **d)** Scatterplot showing changes to 22G-RNAs in *wago-1* as compared to wild type in three different life stages, marked for RIP-Seq targets of ERGO-1 or ALG-3/4 as defined by Seroussi et al. (2023). Overlapping genes and targets of neither are shown in grey. Each dot represents one gene, which typically has several 22G-RNAs mapping to it. **e)** Scatterplot showing changes to mRNA levels in *wago-1* as compared to wild type in three different life stages, marked for RIP-Seq targets of ERGO-1 or ALG-3/4 as defined by Seroussi et al. (2023). Overlapping genes and targets of neither are shown in grey. **f)** Scatterplot showing changes to mRNA levels in *wago-1* as compared to wild type in three different life stages. Argonaute genes are indicated in purple; any with statistically significant changes to mRNA levels are labelled.

The group of 22G-RNA with increased levels in *wago-1* mutants displayed a profile of 3’-end enrichment (Figure 5a), which is the signature of several Argonautes including WAGO-3 and WAGO-9, with which WAGO-1 shares many targets (Seroussi et al., 2023; Tyc et al., 2017). WAGO-3 class 22G-RNAs were changed in both directions (Figure 5c), but WAGO-9 class 22G-RNAs tended to be increased as a group in the gravid adult (Figure 5c). Also in the gravid adult, we saw a general upregulation of WAGO-9 and WAGO-3 target mRNAs, with less effect on the targets shared between them (Supplementary Figure 8b). Possibly, WAGO-9 can partially compensate for loss of WAGO-1. We did not detect strong changes in the 22G-RNAs downstream of the two 26G-RNA pathways, although the subset of highest expression 22G-RNAs downstream of ERGO-1 again tended to be elevated in the L4 (Figure 5d), suggesting a repressive effect of WAGO-1 on the ERGO-1 pathway during spermatogenesis (where it is typically not active).

As mentioned before, at the mRNA level, the most striking effect was observed in gravid adults, where we detected close to 1,400 genes with elevated mRNA levels (Figure 4b). Strikingly, these genes were heavily enriched for ALG-3/4 targets (Figure 5e). The mRNA levels of the two Argonautes, ALG-3/4, themselves were also increased in gravid adult *wago-1* worms along with mRNA levels of WAGO-10, which has been implicated in the same pathway (Figure 5f) (Charlesworth et al., 2021; Seroussi et al., 2023). These changes to ALG-3/4 target mRNAs were specific to the gravid adult stage, with no effect in L4 larvae, where ALG-3/4 are normally expressed, or in embryos (Figure 5e-f). As these increased mRNA changes are not accompanied with lower small RNA changes (Figure 5d) we interpret these changes in mRNA levels as a developmental effect, not as a direct WAGO-1-targeting effect. In line with the increase of ALG-3/4 target mRNAs that we just described, the GO-terms we found for genes upregulated in *wago-1* gravid adult animals were related to sperm development (Supplementary Figure 8c). Next to these GO terms, there was also upregulation of genes related to defence and immunity (Supplementary Figure 8c) as was the case for *wago-3* (Supplementary Figure 5b). While there were fewer significant GO-terms for the genes downregulated in *wago-1* gravid adult animals, we did find that these terms related to cuticle development, molting cycle, and post-embryonic morphogenesis (Supplementary Figure 8d), suggesting mis-regulation of the cuticle and its developmental cycle, though not enough to cause a visible phenotype.

## Discussion

### 22G-mediated mRNA silencing is impossible to predict from *wago* mutant data

One thing that our analyses have consistently shown, is that changes to 22G-RNAs and changes to mRNAs do not globally coincide. While our data does not show why this is the case, it appears that many 22G-RNAs are not effectively driving gene repression. While 22G-RNAs are specific to nematodes, this phenomenon may not be. A study published as a preprint has indicated that mouse pachytene piRNAs only regulate the steady state of a small subset of the more than hundred mRNAs which they cleave (Cecchini et al., 2024). While there are several studies showing examples of small RNAs regulating mRNA levels of specific genes (Barucci et al., 2020; Montgomery et al., 2021; Reed et al., 2019; Rogers & Phillips, 2020), we are not aware of studies that have proven such regulation to occur globally for any small RNA species, and the phenomenon that only a few small RNAs carry significance for mRNA expression may be more common than previously thought.

Nevertheless, a number of potential direct targets may be identifiable through our data. For WAGO-3, we were able to identify a total of 18 potential target genes, where 22G-RNAs were lost and mRNAs were gained in absence of a functional *wago-3*, although five of these were not significantly enriched in RIP-Seq studies (Figure 2c). These 18 genes are interesting candidates for further studies, especially the five genes (*ndx-8*, *F15D4.5*, *F15D4.6*, *C38D9.2*, and *C38D9.13*) which overlapped between L4 larvae and gravid adults. Also of note is the gene *aagr-1*, whose increase of mRNAs in L4 larvae (Figure 2c) we found to be accompanied by a gain in protein expression, based on mass spectrometry (Figure 2d). As we do not currently know what causes some 22G-RNAs to be silencing and others to not take effect, studying these genes further may give more insights into this question.

### Life stage matters

Our analyses further emphasize a previously often overlooked aspect of RNAi-related gene silencing: life stage. Here, we have studied the same mutant animals in different life stages and shown that significant differences exist between different developmental stages. For instance, WAGO-3 appears to affect primarily the embryo, and to prevent the expression of a maternal small RNA pathway during spermatogenesis. WAGO-1 on the other hand has a much stronger effect on the adult hermaphrodite, where it represses spermatogenic gene expression. These effects can convolute our insights based on mutant analyses and RIP-Seq experiments. For instance, possibly WAGO-1 targets key spermatogenesis genes. RIP-Seq studies suggest that WAGO-1 binds more 22G-RNAs against oogenic or constitutive germline genes than spermatogenic ones (Seroussi et al., 2023), but as our results shows a link to the ALG-3/4 pathway, and also that not all 22G-RNAs cause silencing, it may still cause specific silencing of spermatogenesis genes. At the same time, this will result in the loss of 22G-RNAs against these genes, since to make 22G-RNAs, their targets need to be expressed. Hence, in a wild-type ovary, this signature will be invisible. Similarly, relevant gene signatures may be hidden for other WAGO proteins. We note that recently a strong developmental effect was noticed for WAGO-12 (NRDE-3), which was shown to bind CSR-1-like 22G-RNAs in the ovary, but ERGO-1 pathway 22G-RNAs in the somatic cells of the embryo (Chen & Phillips, 2025). To unravel these aspects of the worm small RNA pathways, more life-stage specific studies will be needed using more *wago* mutants and RIP-seq studies from selected mutant backgrounds.

### WAGO-3 affects embryonic gene expression and connects to ERGO-1-interacting WAGOs

As just discussed, *wago-1* and *wago-3* mutants showed impacts in different life stages. For WAGO-3, the effect on mRNA level was the strongest in the embryo (Figure 2b). This fits with the current knowledge of WAGO-3, namely that it plays a role in inheritance (Schreier et al., 2025). While WAGO-3 does affect other life stages, it appears that the major role of WAGO-3 is in establishing the inheritance response in the embryo. Interestingly, our data also shows that the system can re-equilibrate later in development, or else stark changes to mRNA levels should have persisted in our L4 larvae and gravid adult samples. Perhaps the Argonaute WAGO-6 plays a role in this, as WAGO-6 is overexpressed in *wago-3* mutant embryos (Figure 3g). Both WAGO-6 class 22G-RNAs and WAGO-8/12 class 22G-RNAs are enriched in the *wago-3* mutants (Figure 3d-e), though the increase of WAGO-6 class 22G-RNAs is less striking in the embryo (Figure 3d), where fewer genes were affected at 22G-RNA level (Figure 2a). It is possible that WAGO-6 gets activated in the embryo when WAGO-3 is missing, and that this activation leads to a general activation of the WAGO-6/8/12 pathway at later life stages as well. Curiously, WAGO-6, −8 and −12 have all been implicated in the ERGO-1 pathway (Seroussi et al., 2023) and we detected an increase of ERGO-1 pathway 22G-RNAs in the L4 larvae (Figure 3f). This suggests a possible connection between WAGO-3 and the maternal (ERGO-1) 26G-RNA pathway.

These findings are supported by our observations of the changing 22G-RNA metagene profiles that accompanied loss of WAGO-3 (see Results; Figure 3a-c). The observation that different WAGOs have different metagene profiles is an interesting one, and the differences we see in these profiles between genes with depleted or elevated 22G-RNA levels can be quite striking (Figure 3c). Nonetheless, while this is a useful tool for assessing trends, many Argonautes share profiles, and the effects observed are likely a result of mixed effects stemming from several Argonautes, and one should be weary of drawing too strong conclusions based on metagene profiling alone, without other supporting data.

### WAGO-1 is connected to silencing of the paternal 26G-RNA pathway during oogenesis

Similar to the connection between WAGO-3 and the maternal ERGO-1 pathway during spermatogenesis, we also detected a connection between WAGO-1 and the paternal (ALG-3/4) 26G-RNA pathway during oogenesis. This suggests that WAGO-1 and WAGO-3 have somewhat complementary effects, both playing a role in repressing 26G-RNA pathways. Interestingly, a recent preprint confirms these results, showing that a different *wago-1* mutant allele than the one used here (*wago(tm1414))* had similar effects; 22G-RNAs against ALG-3/4 targets were strongly increased in adult animals (DiNardo et al., 2025). The connection to the ALG-3/4 pathway is further supported by the increase of spermatogenic genes in the adult worm (Supplementary Figure 8c). Perhaps WAGO-1 is responsible for silencing the ALG-3/4 pathway past spermatogenesis, although the fact that ALG-3/4 target 22G-RNAs were unchanged would suggest that this could also be a secondary effect. The same is the case for WAGO-9. We saw a strong increase of WAGO-9 target mRNAs, which could suggest linkage between WAGO-9 and the ALG-3/4 pathway. While we cannot rule out that WAGO-9 compensates for WAGO-1 (we saw an increase of WAGO-9 target 22G-RNAs) it is also possible that WAGO-1 does not interact directly with WAGO-9: disturbances caused by loss of WAGO-1 may enhance production of WAGO-9 target mRNAs which in turn may cause an increase of WAGO-9 target 22G-RNAs due to higher availability.

### Final thoughts

This study highlights several links between different Argonaute pathways. One interesting observation is that loss of WAGO-3 affects WAGO-6 and possibly also WAGO-8. Curiously, WAGO-6 and WAGO-8 are somatic, while WAGO-3 is expressed in the germline (Seroussi et al., 2023). As we see enrichment of WAGO-6 mRNAs in *wago-3* embryo samples (Figure 3g), it is possible that paternal/maternal WAGO-3 interacts with a nuclear Argonaute (possibly WAGO-12(NRDE-3), Figure 3e; or WAGO-9(HRDE-1) (Schreier et al 2025)) to suppress expression of WAGO-6 during early embryogenesis, leading to transcriptionally repressed *wago-6* also in the somatic cells. Alternatively, WAGO-6 may be ectopically expressed in the germline in *wago-3* mutants.

Another interesting observation is that loss of either WAGO-1 or WAGO-3, did not cause loss of 22G-RNAs against their cognate target genes (Supplementary Figure 3c-d and Supplementary Figure 7c-d). We have recently shown the same for mutants lacking the closely related WAGO, WAGO-4 (Seistrup et al., 2025). Currently, the general consensus in the field remains that such observations are caused by complementation by other Argonautes. Our study supports that idea, as we describe evidence for shifts of 22G-RNA populations between WAGO proteins.

Furthermore, our work highlights that the mere presence of 22G RNAs against a particular gene does not automatically imply effective silencing of that gene. The reasons behind this observation are not clear, but it does show that claiming any biological effect of a specific WAGO-target relationships requires more than simply 22G RNA matching.

Finally, all Argonaute targets mentioned here have been determined via RIP-Seq using endogenously, N-terminally tagged versions of the proteins. While animals carrying such tags appears phenotypically wild type, we cannot exclude that these tags may play a role in 22G-RNA selection and/or binding.

## Methods

### C. elegans strains and harvest

Worms were cultured on nematode growth medium (NGM) plates (90 mm diameter) seeded with Escherichia coli OP50 in temperature-controlled incubators at 20°C. Worms were synchronized for life stage via bleaching and grown on egg plates, the preparation of which is described below. Worms were bleached a second time before being transferred to standard NGM plates, in the case of harvesting of L4 larvae or adult worms, or to egg plates when harvesting embryos. Harvesting was carried out using M9 buffer and aliquots were fast-frozen on dry ice before storage at −80°C.

Egg plates were made by mixing egg yolk with 50 ml LB medium per egg. After incubation at 65 °C for 2–3 hours, the mixture was cooled to room temperature before adding 10 ml OP50 culture per egg. About 10 ml was added to standard NGM plates (90 mm diameter) and incubated at room temperature. The next day, excess liquid was removed, and egg plates were incubated at room temperature for another 2 days.

Deletion mutants were confirmed via PCR using Taq polymerase (New England Biolabs) using the standard protocol and an annealing temperature of 60 °C for all genotypes; primers specified below. All deletion mutants were outcrossed to N2 wild type males. Heterozygous offspring thereof were crossed to N2 wild type hermaphrodites and offspring were kept homozygous for the deletion for five generations before harvest.

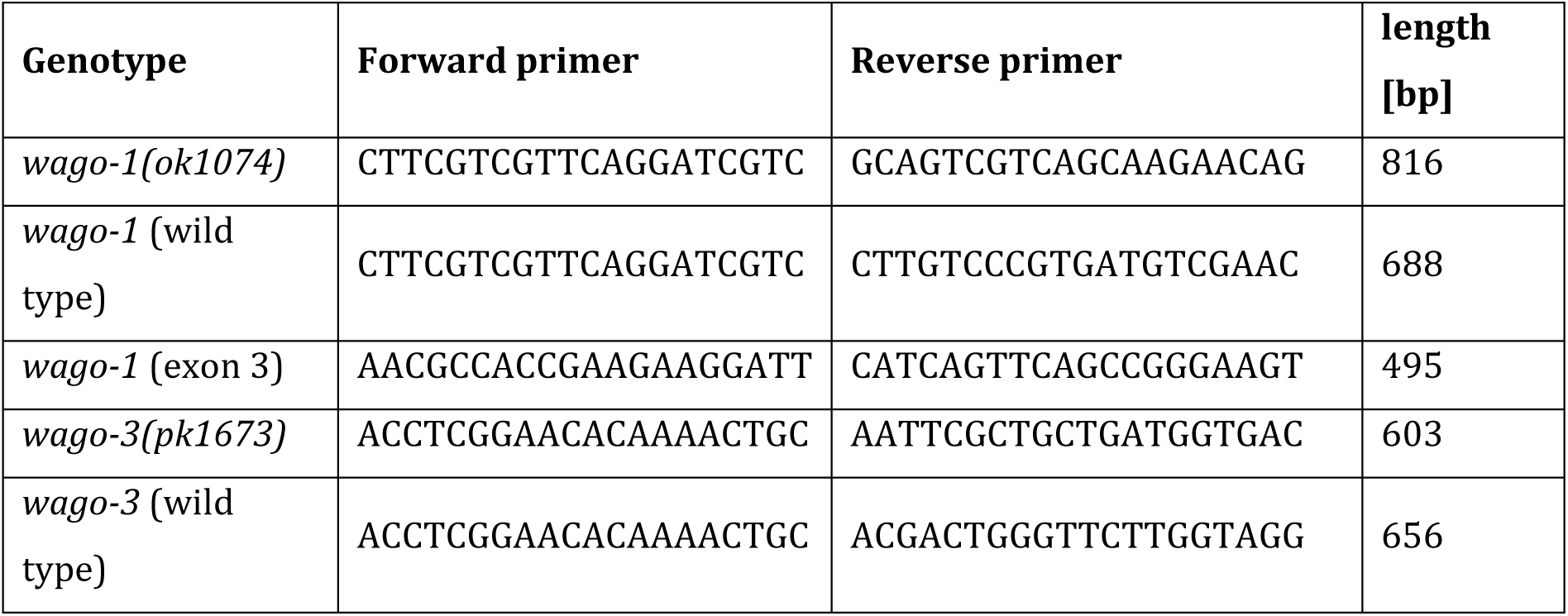

### Immunoprecipitation

Previously collected aliquots of ∼300 μL L4 larvae were thawed on ice and mixed with the same volume of 2× lysis buffer (50 mM Tris HCl pH 7.5, 300 mM NaCl, 3 mM MgCl2, 2 mM dithiothreitol (DTT), 0.2% Triton X-100, cOmplete Mini EDTA-free Protease Inhibitor Cocktail (11836170001, Roche)). For mass spectrometry analysis, samples were sonicated using a Bioruptor Plus device (B01020001, Diagenode) (4°C, 10 cycles, 30 seconds on and 30 seconds off). For sequencing, samples were dropped into liquid nitrogen in a mortar, forming little balls that were subsequently pulverised using a pestle. The powder from this was transferred to a douncer, where it was dounced 40 times before turning liquid.

For both types of experiments, samples were next centrifugated for 10 min at 4°C and 21,000*g* and supernatants were transferred into a fresh tube. Total protein concentrations of soluble worm extracts were determined using the Pierce BCA Protein-Assay (23225, Thermo Fisher Scientific) and an Infinite M200 Pro plate reader (Tecan), and extracts were diluted with 1× lysis buffer to reach similar concentrations and a volume of 500 μL in the case of mass spectrometry samples and 550 μL for the preparation of sequencing samples. In the latter case, 50 μL were removed to serve as input control.

For each experiment, 30 μl Novex DYNAL Dynabeads Protein G (10004D, Invitrogen) were washed three times with 500 μl 1× wash buffer (25 mM Tris HCl pH 7.5, 150 mM NaCl, 1.5 mM MgCl2, 1 mM DTT, cOmplete Mini EDTA-free Protease Inhibitor Cocktail), combined with the remaining 500 μl extract and incubated with rotation for 1 hour at 4°C. In the meantime, 8 μg antibody (monoclonal anti-Flag M2, F3165, Sigma-Aldrich) was conjugated to another 30 μl Novex DYNAL Dynabeads Protein G according to the manufacturer’s instructions. Extracts were separated from non-conjugated Dynabeads, combined with antibody-conjugated Dynabeads and incubated with rotation for 2 hours at 4 °C. Following three washes with 500 μl 1× wash buffer, antibody-conjugated Dynabeads were resuspended in 50 μl nuclease-free water. For mass spectrometry analyses, extracts were instead resuspended in 25 μl 1.2× Novex NuPAGE LDS sample buffer (NP0007, Invitrogen) supplemented with 120 mM DTT and incubated at 70 °C for 10 minutes before fast-freezing in dry ice and storage at −80°C.

### Mass spectrometry

Samples were prepared in quadruplicates and thawed on ice before being separated on a Novex NuPAGE 4–12% Bis-Tris Mini Protein Gel (NP0321, Invitrogen) in 1× Novex NuPAGE MOPS SDS Running Buffer (NP0001, Invitrogen) at 180 V for 10 minutes. After separation, the samples were processed by in-gel digest as described in (Shevchenko et al., 2006). After protein digest, the peptides were desalted using a C18 StageTip78. For measurement, the digested peptides were separated on a 25 cm reverse-phase capillary (inner diameter, 75 μm) packed with Reprosil C18 material (Dr. Maisch). Elution was carried out along a 2 hour gradient of 2–40% of a mixture of 80% acetonitrile/0.5% formic acid using the EASY-nLC 1000 system (LC120, Thermo Fisher Scientific). A Q Exactive Plus mass spectrometer (Thermo Fisher Scientific) operated with a Top10 data-dependent MS/MS acquisition method per full scan was used for measurement. Processing of the obtained results was performed with the MaxQuant software (v.1.5.2.8) against the Wormbase protein database (version WS270) for quantification.

### RNA extraction

Harvested frozen worms were thawed on ice and 400 µl TRIzol LS Reagenz (10296010, Invitrogen) was added to the sample. The worm samples were then lysed by six cycles of freezing in liquid nitrogen followed by thawing at 37°C and vortexing. Embryo samples were lysed by being dropped into liquid nitrogen in a mortar, forming little balls that were subsequently pulverised using a pestle. Then 400 µl TRIzol LS Reagenz (10296010, Invitrogen) was added to the embryo samples. For all samples, liquid was transferred to a new tube and centrifuged for 5 min at 4°C and 21,000g. Supernatants were transferred into a fresh tube and RNA was extracted using Direct-Zol RNA Miniprep Kit (R2052, Zymo Research) according to the manufacturer’s instructions and resuspended in nuclease-free water. RNA quality and quantity was assessed initially using the Bioanalyzer RNA 6000 Nano Kit (5067-1511, Agilent Technologies) and subsequently using the Qubit RNA BR Assay Kit (Q10210, Invitrogen). The same extracted RNA was used for sequencing of small RNAs and of mRNAs.

### Small RNA library preparation and sequencing

NGS library prep was performed with NEXTflex Small RNA-Seq Kit V3 following Step A to Step G of Bioo Scientific’s standard protocol (V19.01) with a ligation of the 3’ 4N Adenylated Adapter over night at 20°C. Previously to library prep, all samples were treated with RNA 5’ Pyrophosphohydrolase (RppH, NEB). Libraries were prepared with a starting amount of 360 ng, addition of 1 µl NEXTflex tRNA/YRNA Blockers (BiooScientific) and amplified in 15 PCR cycles. Amplified libraries were purified by running an 8% TBE gel and size-selected for 144 – 163 nt. Libraries were profiled in a High Sensitivity DNA on a 2100 Bioanalyzer (Agilent technologies) and quantified using the Qubit dsDNA HS Assay Kit, in a Qubit 2.0 Fluorometer (Life technologies). All samples were pooled in equimolar ratio and sequenced on a NextSeq 500/550 Flowcell, SR for 1x 75 cycles plus 6 or 7 cycles for the index read

### mRNA library preparation and sequencing

NGS library prep was performed with Illumina’s Stranded mRNA Prep Ligation Kit following Stranded mRNA Prep Ligation ReferenceGuide (June 2020) (Document # 1000000124518 v00). Libraries were prepared with a starting amount of 500 ng and amplified in 10 PCR cycles. For normalization, 1 µl of a 1:100 dilution of ERCC spike-ins (Ambion) was added. Libraries were profiled in a High Sensitivity DNA on a 2100 Bioanalyzer (Agilent technologies) and quantified using the Qubit dsDNA HS Assay Kit, in a Qubit 2.0 Fluorometer (Life technologies). All 23 samples were pooled in equimolar ratio and sequenced on a NextSeq500 Highoutput FC, SR for 1x 80 or 1×79 cycles plus 10 cycles for the index read and 1 or 2 dark cycle upfront.

### Read processing and mapping

Raw sequenced reads from high-quality libraries, as assessed by FastQC, were processed using Cutadapt (Martin, 2011) for adapter removal (-a TGGAATTCTCGGGTGCCAAGG -O 5 -m 26 -M 48) and low-quality reads were filtered out using the FASTX-Toolkit (fastq_quality_filter, -q 20 -p 100 -Q 33). Unique molecule identifiers were used to remove PCR duplicates using a custom script and were subsequently removed using seqtk (trimfq-l 4 – b 4). Finally, in the case of small RNA-Seq, reads shorter than 15 nucleotides were removed using seqtk (seq -L 15). A custom GTF file was created by adding transposons retrieved from Wormbase (PRJNA13758 .WS264) to the Ensembl reference WBcel235.84 and reads were aligned using bowtie (v.1.3.1) (Langmead et al., 2009)(--phred33-quals --tryhard --best --strata --chunkmbs 256 -v 2 -M 1) for small RNA-Seq or subread v.2.0.0 (Liao et al., 2013) featureCounts (--donotsort -t exon) for mRNA-Seq.

For sRNA-Seq, length distributions and 5’-nucleotides of unmapped files were determined with a Python script available at https://github.com/Tunphie/SequencingTools/blob/main/summarizeNucleotideByReadLength.py. Small RNA types from mapped libraries were classified using a Python script available at https://github.com/Tunphie/SequencingTools/blob/main/smRNA_TypeCounter.py, using strict classification rules, before 22G-RNAs were extracted using a python script available at https://github.com/adomingues/filterReads with less strict rules: they were defined as any read of length 20-23 nt with no bias at the 5’-position and no requirement of mapping orientation. These defined 22G-RNAs were used for further analysis for both RIP-Seq and for small RNA sequencing of the deletion mutants.

### Coverage tracks

BigWig files were generated from mapped mRNA reads using deepTools v.3.5.5 bamCoverage (Ramírez et al., 2016) using default settings. These were visualised in the online tool Integrative Genomics Viewer (IGV)(Robinson et al., 2011). Tracks were downloaded as single vector graphics (SVG) files and recoloured using Adobe Illustrator.

### Differential analysis

Reads were counted using htseq-count (v.2.0.2) (Anders et al., 2015) (-s no -m intersection-nonempty). Reads per kb of transcript per million mapped reads (RPKM) values were calculated relative to all mapped reads.

For RIP-Seq experiments, targets were defined as genes that were positive in at least two out of three replicates, with positive meaning that the 22G-RNA RPKM was (1) above 4 in the IP; (2) at least twice as high in the IP relative to the input; and (3) non-zero in the input. For 22G-RNAs mapping to transposons, RPKM values were calculated relative to the predicted number and length of insertions in the custom annotation file and positives were defined using only requirements 2 and 3 above with no minimal RPKM requirement in the IP.

For deletion mutant experiments, targets were determined using DeSeq2 (Love et al., 2014) in R, with a p-value cutoff of 0.01 and a cutoff for log2(fold change) of 1. For mRNA-Seq, reads mapping to ERCCs were used for normalization by resetting the size factors using estimateSizeFactors() on the data with only ERCC reads. All data was shrunk using lfcShrink() using the method ashr (Stephens, 2016) Our data was compared to published RIP-Seq data from (Schreier et al., 2022), in the case of WAGO-3, and (Seroussi et al., 2023) for all other Argonautes.

Comparisons were carried out in R using the packages ggplot2 or UpsetR (Conway et al., 2017). GO-term analyses were carried out in R using enrichGO() from clusterProfiler (Yu et al., 2012) using default settings.

## Strains and Reagents

| Strain | Acquisition |
| --- | --- |
| N2 Bristol isolate (wild type) | *Caenorhabditis* Genetics Center (CGC) |
| *gfp::3xflag::wago-3(xf119)* | (Schreier et al., 2022) |
| *wago-1(ok1074)* | *Caenorhabditis* Genetics Center (CGC) |
| *wago-3(pk1673)* | *Caenorhabditis* Genetics Center (CGC) |

| Chemical/Reagent | Company |
| --- | --- |
| TRIzol LS Reagenz | Invitrogen |
| Direct-Zol RNA Miniprep Kit | Zymo Research |
| NEXTflex Small RNA-Seq Kit V3 | BiooScientific |
| RNA 5' Pyrophosphohydrolase (RppH) | New England Biolabs (NEB) |
| NEXTflex tRNA/YRNA Blockers | BiooScientific |
| Illumina's Stranded mRNA Prep Ligation Kit | Illumina |
| Taq polymerase | New England Biolabs (NEB) |
| cOmplete Mini EDTA-free Protease Inhibitor Cocktail | Roche |
| Pierce BCA Protein-Assay | Thermo Fisher Scientific |
| Novex DYNAL Dynabeads Protein G | Invitrogen |
| Monoclonal anti-Flag M2 antibody | Sigma-Aldrich |
| Novex NuPAGE LDS sample buffer | Invitrogen |
| Novex NuPAGE MOPS SDS Running Buffer | Invitrogen |
| ReproSil-Pur Basic-C18 | Dr. Maisch |

## Data Availability

RIP-Seq data has been uploaded to the European Nucleotide Archive (ENA, https://www.ebi.ac.uk/) under the accession number PRJEB102712. All other sequencing data has been uploaded to ENA under the accession number PRJEB95105.

The mass spectrometry proteomics data have been deposited to the ProteomeXchange Consortium via the PRIDE (Perez-Riverol et al., 2025) partner repository with the dataset identifier PXD072057.^1^

## Acknowledgements

The authors thank Jan Schreier, Ida Josefine Isolehto, and Svenja Hellmann for their help in the lab as well as the entire Ketting lab for their inputs during discussions. We thank the genomics core facility at the IMB for their assistance in library preparation.

---

**Supplementary Figure 1:**
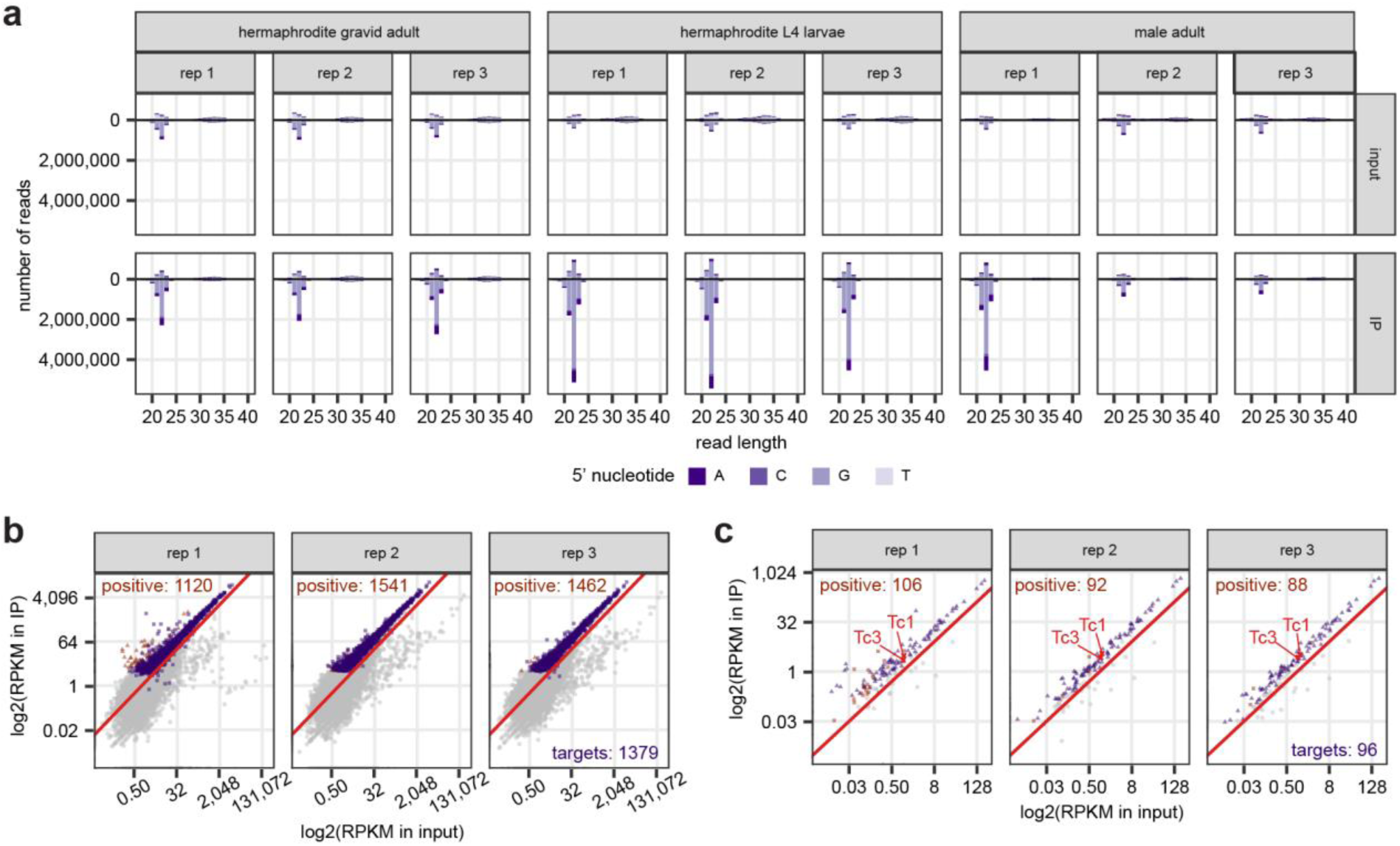
RNA-immunoprecipitation sequencing of GFP::3xFLAG::WAGO-3(*xf119*). **a)** Bar plot showing the distribution of reads from our sequencing libraries of L4 hermaphroditic larvae as well as libraries from male and hermaphrodite adult animals from Schreier et al., 2022. Colours represent the 5’-nucleotide of each read. **b)** Scatterplot showing 22G-RNA reads mapping to different genes excluding transposons. Positive hits in each replicate are indicated in orange and targets (positive in two or more replicates) are indicated in purple. Each dot represents one gene which typically had several 22G-RNAs mapped to it. **c)** Scatterplot showing 22G-RNA reads mapping to transposons. Positive hits in each replicate are indicated in orange and targets (positive in two or more replicates) are indicated in purple. Tc1 and Tc3 are highlighted in red. Each dot represents one gene which typically had several 22G-RNAs mapped to it.

**Supplementary Figure 2:**
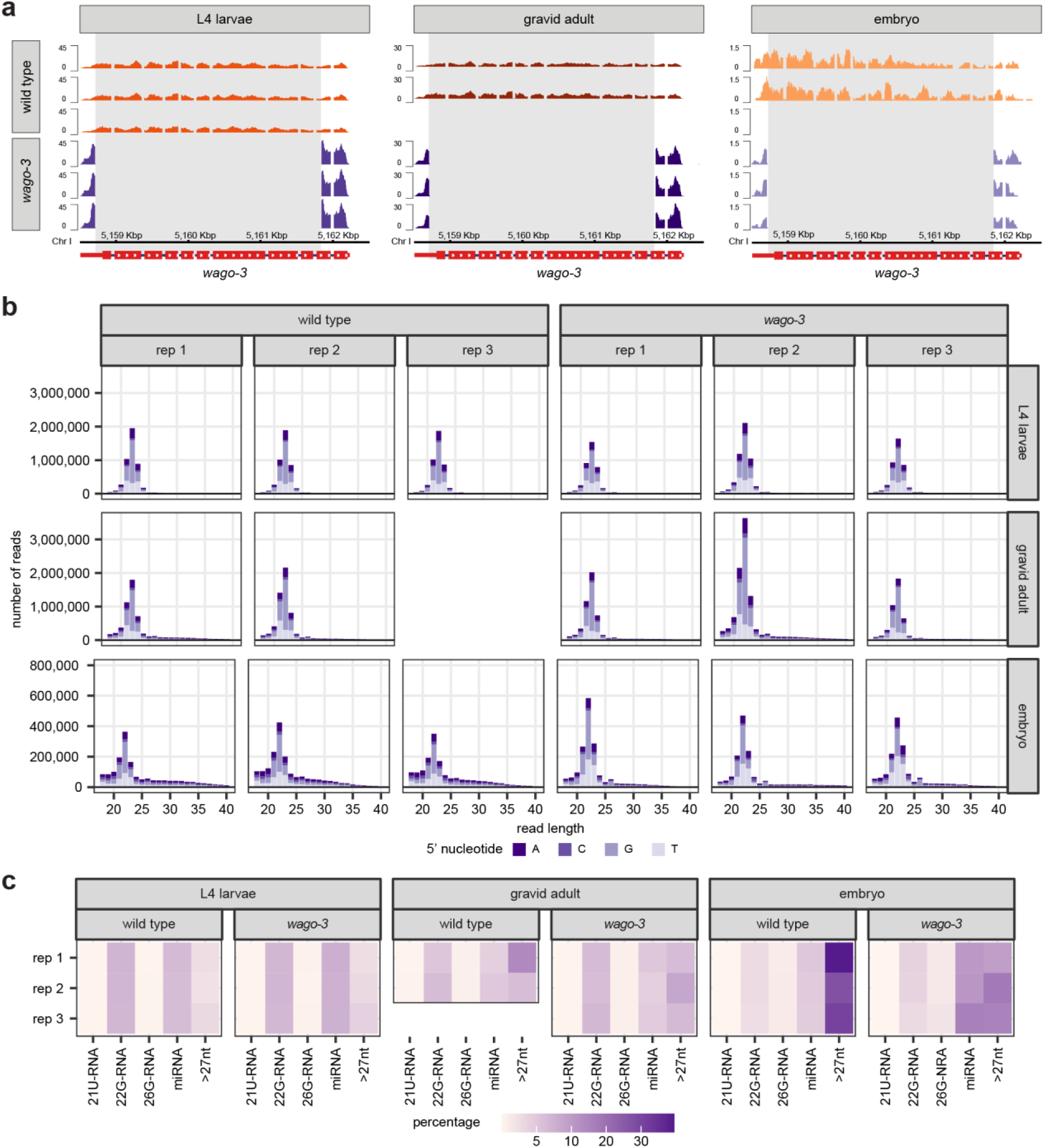
Sequencing profiles of *wago-3* and wild type libraries. **a)** mRNA reads along the *wago-3* gene in wild type and *wago-3* deletion mutants in L4 larvae, gravid adult worms, and embryos. Location of deletion marked as a grey box. **b)** Length distribution profiles of all, unmapped reads from small RNA sequencing libraries from wild type and *wago-3* L4 larvae, gravid adult, and embryo samples. Colours represents 5’-nucleotide. **c)** Percentwise distribution of 21U-RNAs, 22G-RNAs, 26G-RNAs, miRNAs and reads longer than 27 nucleotides in unmapped small RNA sequencing samples from wild type and *wago-3* L4 larvae, gravid adult worms, and embryos.

**Supplementary Figure 3:**
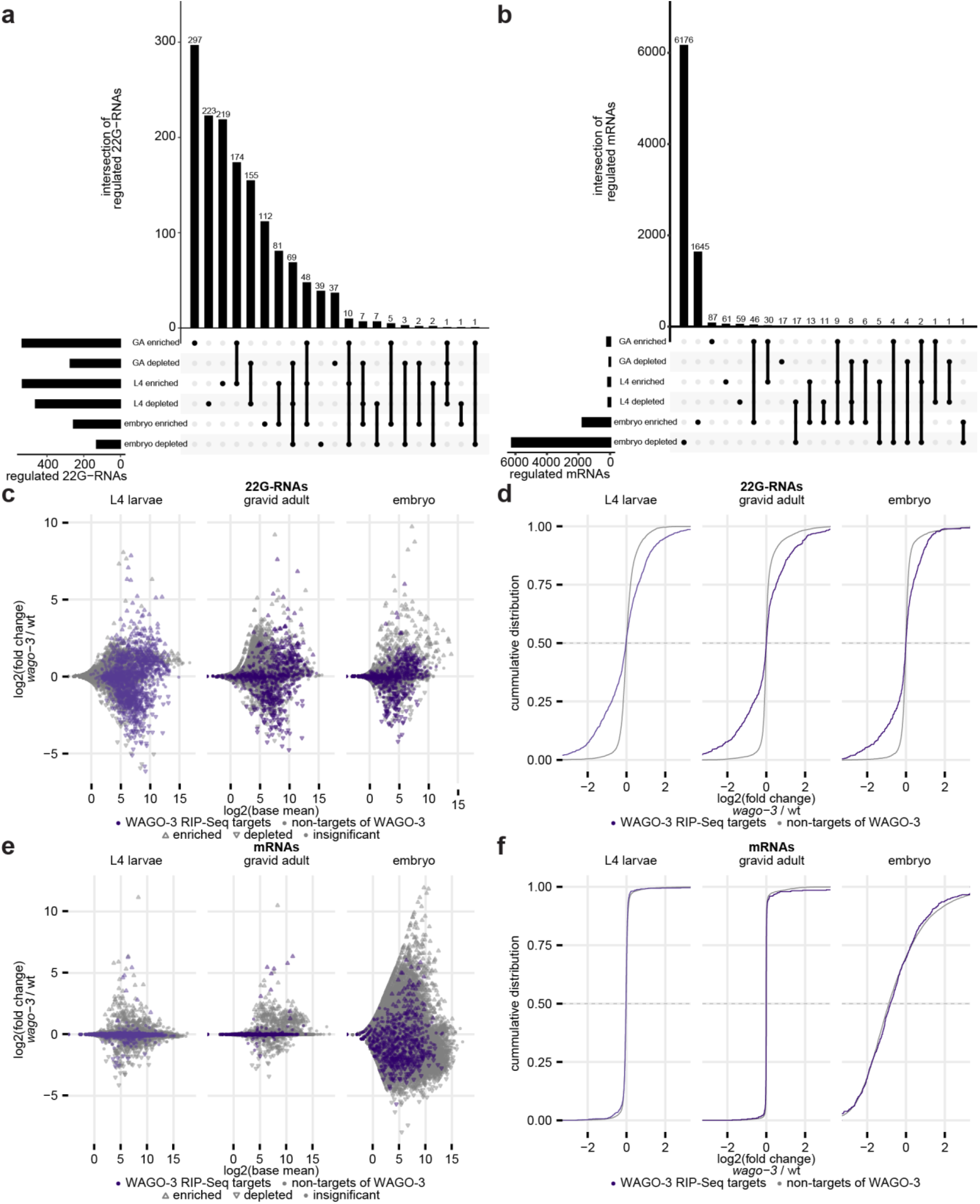
Behaviour of WAGO-3 target genes in *wago-3* mutant animals. **a)** UpSet plot showing overlaps between all genes with statistically significant changes to 22G-RNA levels (enriched or depleted) in *wago-3* as compared to wild type, in L4 larvae (L4), gravid adult (GA), and embryo samples. **b)** UpSet plot showing overlaps between all genes with statistically significant changes to mRNA levels (enriched or depleted) in *wago-3* as compared to wild type, in L4 larvae (L4), gravid adult (GA), and embryo samples. **c)** Scatterplot showing changes to 22G-RNAs in *wago-3* as compared to wild type in three different life stages, marked for WAGO-3 RIP-Seq targets. For L4 larvae, results from RIP-Seq of L4 larvae (Figure 1) was used as the comparison. For gravid adult worms and embryos, results from RIP-Seq of gravid adult worms was used (Schreier et al., 2022). Each dot represents one gene, which typically has several 22G-RNAs mapping to it. **d)** CDF plot quantifying panel c. X-axis has been shortened to show the relevant area. **e)** Scatterplot showing changes to mRNAs in *wago-3* as compared to wild type in three different life stages, marked for WAGO-3 RIP-Seq targets. For L4 larvae, results from RIP-Seq of L4 larvae (Figure 1) was used as the comparison. For gravid adult worms and embryos, results from RIP-Seq of gravid adult worms was used (Schreier et al., 2022). **f)** CDF plot quantifying panel e. X-axis has been shortened to show the relevant area.

**Supplementary Figure 4:**
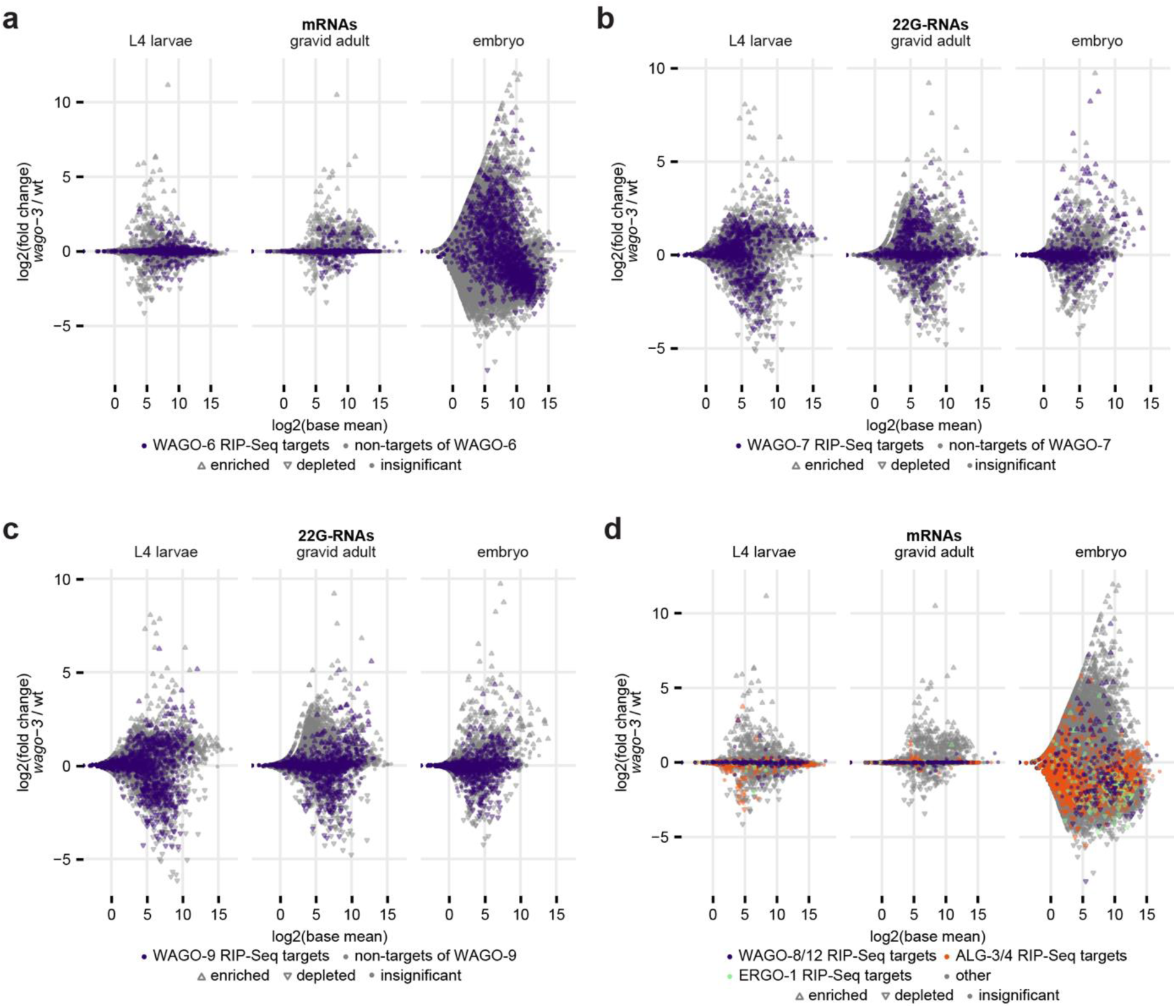
Changes to mRNA levels of ERGO-1-interacting Argonautes in *wago-3*. **a)** Scatterplot showing changes to mRNAs in *wago-3* as compared to wild type in three different life stages, marked for RIP-Seq targets of WAGO-6 as defined by Seroussi et al. (2023). **b)** Scatterplot showing changes to mRNAs in *wago-3* as compared to wild type in three different life stages, marked for RIP-Seq targets of WAGO-7 as defined by Seroussi et al. (2023). **c)** Scatterplot showing changes to mRNAs in *wago-3* as compared to wild type in three different life stages, marked for RIP-Seq targets of WAGO-9 as defined by Seroussi et al. (2023). **d)** Scatterplot showing changes to mRNAs in *wago-3* as compared to wild type in three different life stages, marked for RIP-Seq targets of WAGO-8/12, ERGO-1, or ALG-3/4 as defined by Seroussi et al. (2023). Overlapping genes and targets of neither are shown in grey.

**Supplementary Figure 5:**
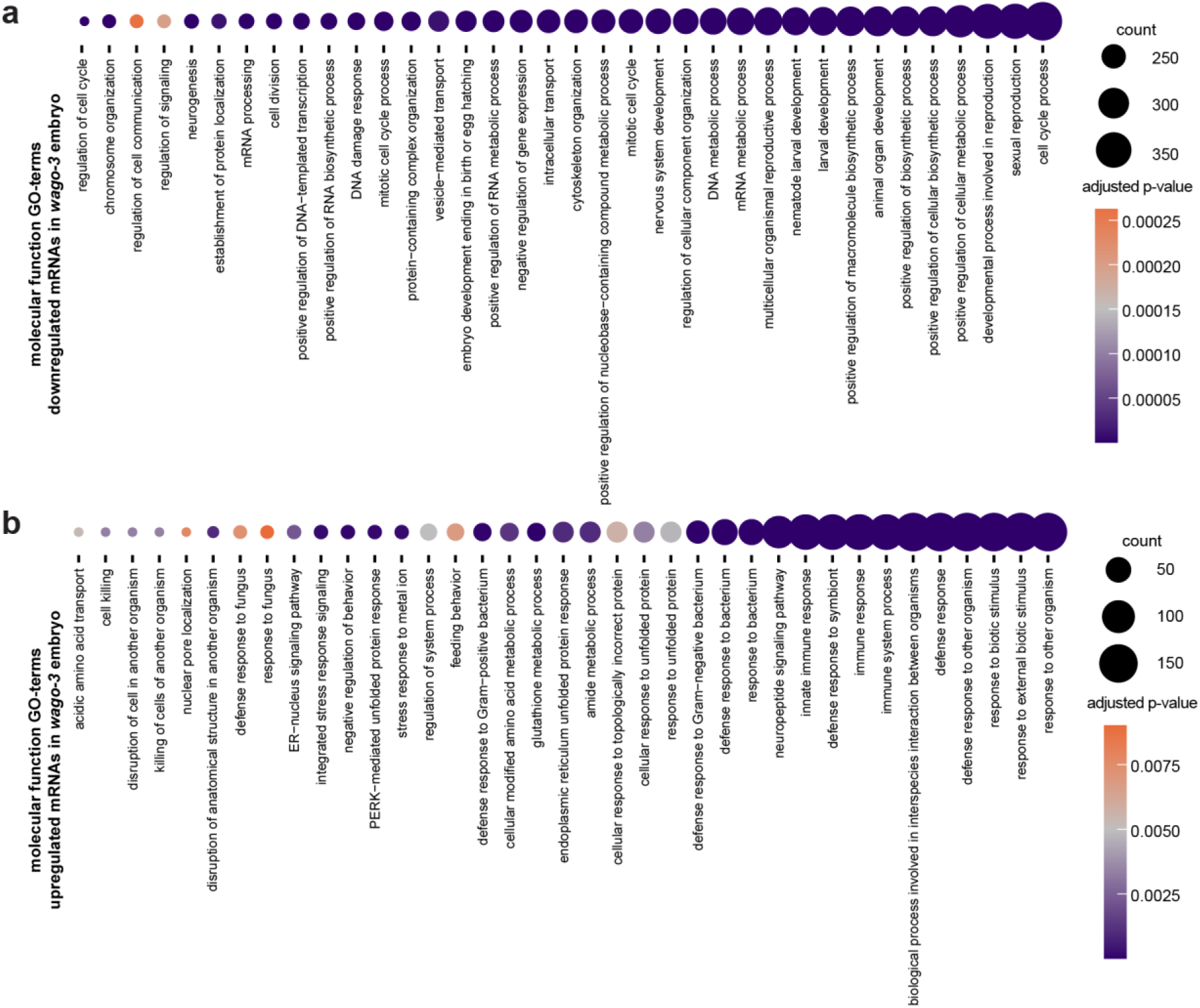
GO-term associated with changing mRNAs in *wago-3* embryos. **a)** Dot plot showing all ‘molecular function’ GO-terms with adjusted p-values < 0.01 and a count of 200 or more related to genes with downregulated mRNAs in *wago-3* embryo libraries. **b)** Dot plot showing all ‘molecular function’ GO-terms with adjusted p-values < 0.01 related to genes with upregulated mRNAs in *wago-3* embryo libraries.

**Supplementary Figure 6:**
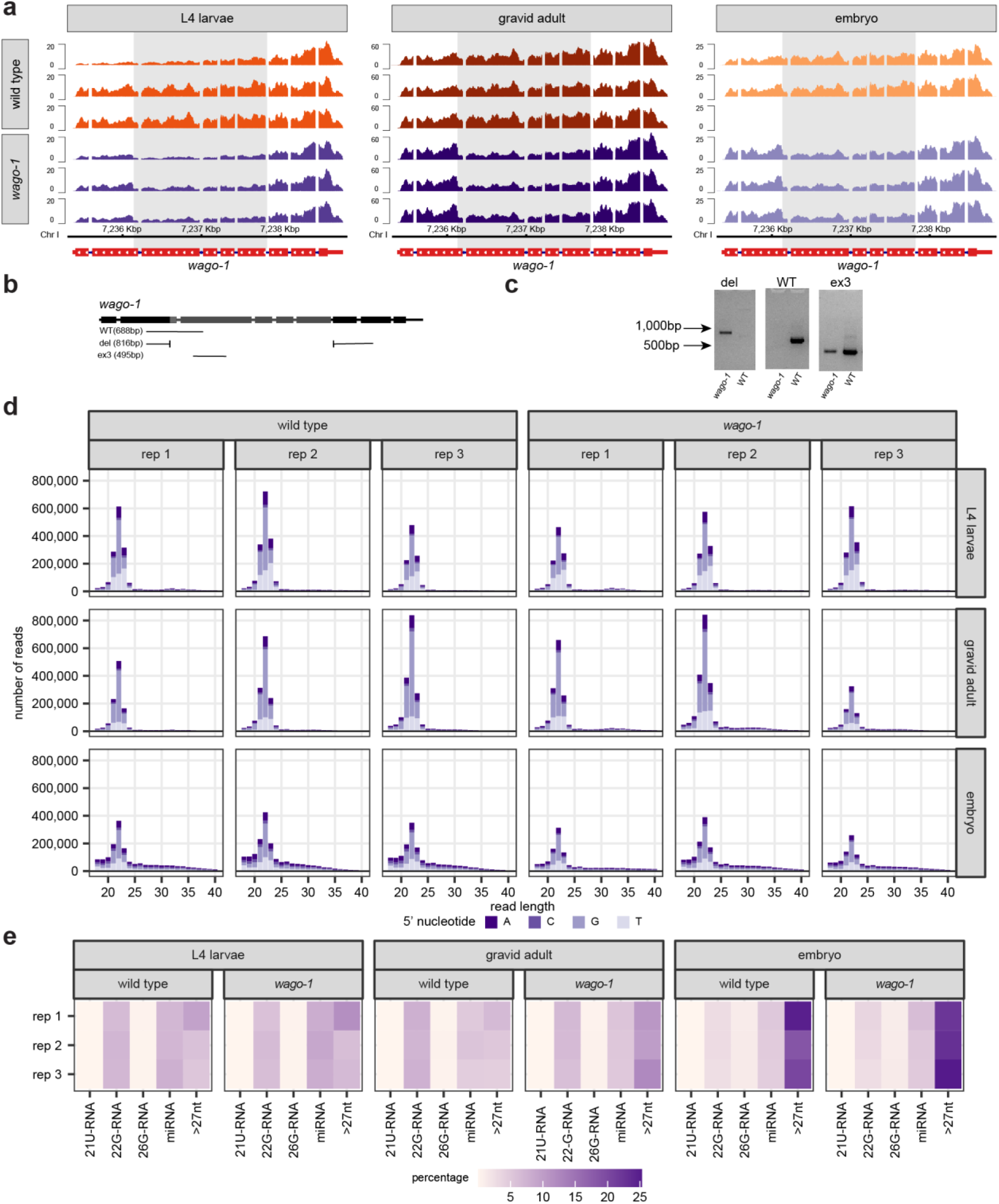
Sequencing profiles of *wago-1* and wild type libraries. **a)** mRNA reads along the *wago-1* gene in wild type and *wago-1* deletion mutants in L4 larvae, gravid adult worms, and embryos. Location of deletion marked as a grey box. **b)** Schematic of the *wago-1* gene and the locations of the primers used for genotyping (WT for wild type band, del for *wago-1* deletion band; spanning the deletion, and ex3 for internal *wago-1* deletion band in exon 3). Presumed deletion marked in grey. **c)** PCR genotyping of wild type and *wago-1* mutant worms using the primers presented in panel b. **d)** Length distribution profiles of all, unmapped reads from small RNA sequencing libraries from wild type and *wago-3* L4 larvae, gravid adult, and embryo samples. Colours represents 5’-nucleotide. **e)** Percentwise distribution of 21U-RNAs, 22G-RNAs, 26G-RNAs, miRNAs and reads longer than 27 nucleotides in unmapped small RNA sequencing samples from wild type and *wago-3* L4 larvae, gravid adult worms, and embryos.

**Supplementary Figure 7:**
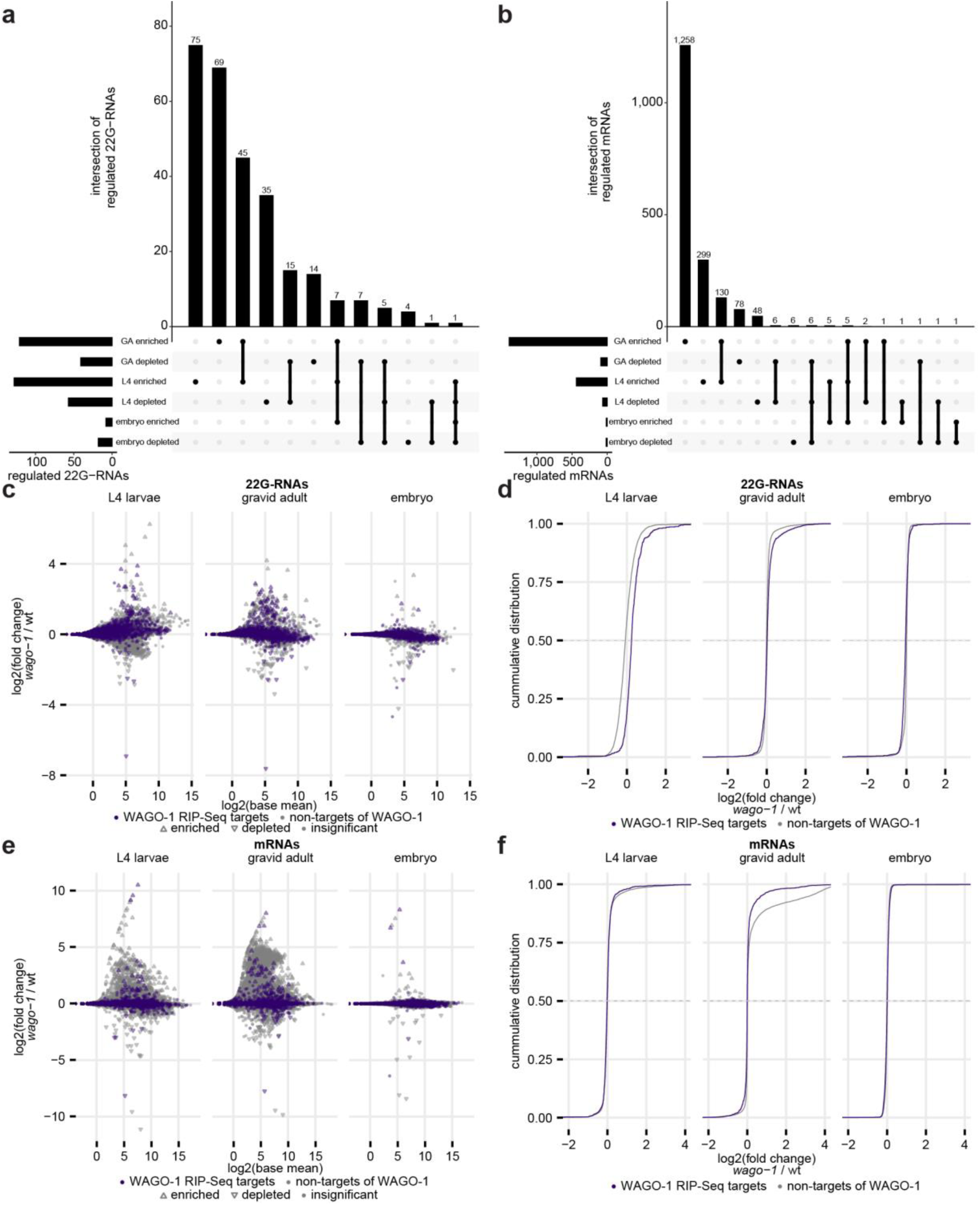
Behaviour of WAGO-1 target genes in *wago-1* mutant animals. **a)** UpSet plot showing overlaps between all genes with statistically significant changes to 22G-RNA levels (enriched or depleted) in *wago-1* as compared to wild type, in L4 larvae (L4), gravid adult (GA), and embryo samples. **b)** UpSet plot showing overlaps between all genes with statistically significant changes to mRNA levels (enriched or depleted) in *wago-1* as compared to wild type, in L4 larvae (L4), gravid adult (GA), and embryo samples. **c)** Scatterplot showing changes to 22G-RNAs in *wago-1* as compared to wild type in three different life stages, marked for WAGO-1 RIP-Seq targets as defined by Seroussi et al. (2023). Each dot represents one gene, which typically has several 22G-RNAs mapping to it. **d)** CDF plot quantifying panel c. X-axis has been shortened to show the relevant area. **e)** Scatterplot showing changes to mRNAs in *wago-1* as compared to wild type in three different life stages, marked for WAGO-1 RIP-Seq targets as defined by Seroussi et al. (2023). **f)** CDF plot quantifying panel e. X-axis has been shortened to show the relevant area.

**Supplementary Figure 8:**
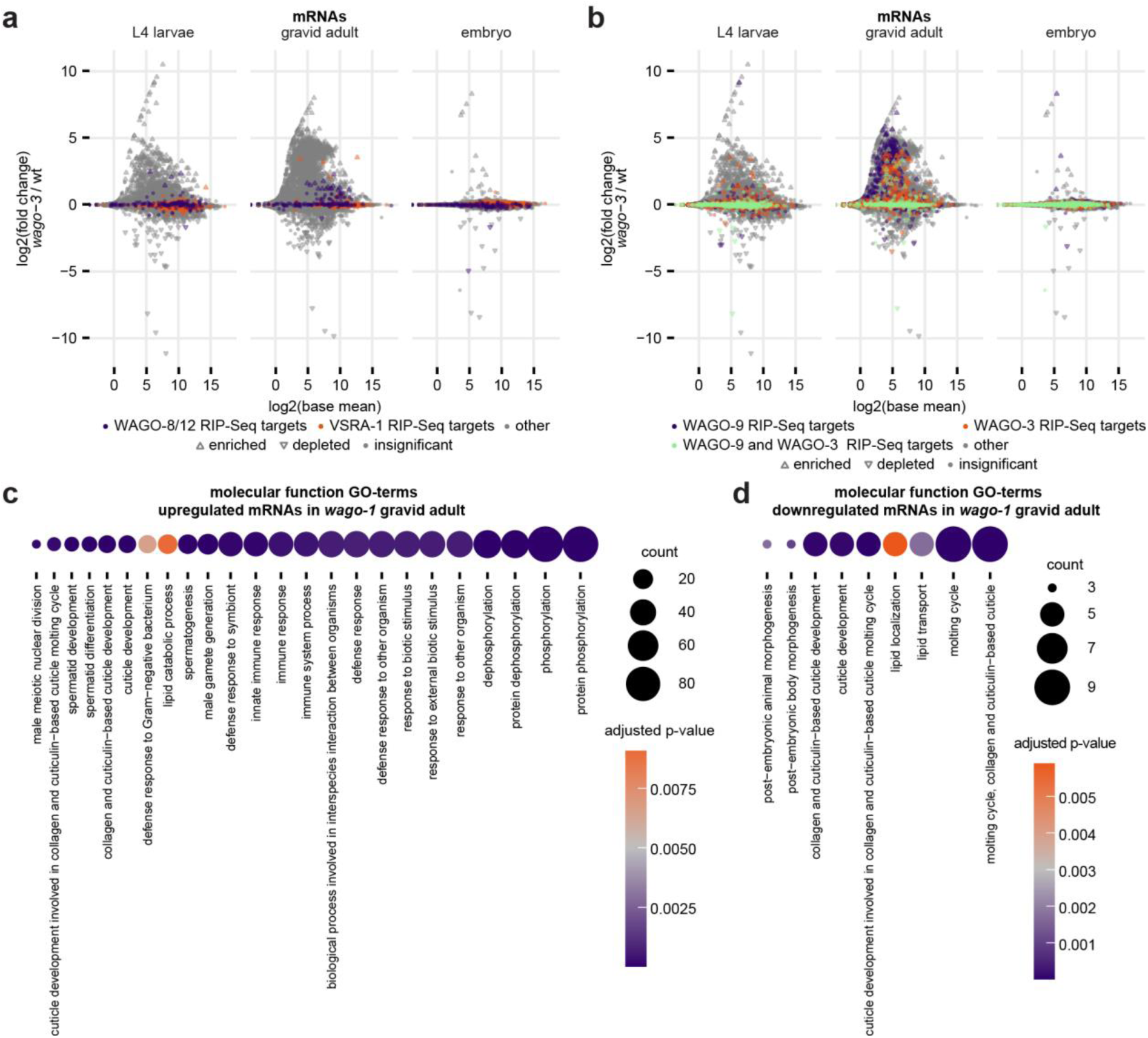
Trends of mRNA changes in *wago-1*. **a)** Scatterplot showing changes to mRNA levels in *wago-1* as compared to wild type in three different life stages, marked for RIP-Seq targets of WAGO-8/12 or VSRA-1 as defined by Seroussi et al. (2023). Overlapping genes and targets of neither are shown in grey. **b)** Scatterplot showing changes to mRNAs in *wago-1* as compared to wild type in three different life stages, marked for RIP-Seq targets of WAGO-9 (Seroussi et al., 2023) and/or WAGO-3 (Schreier et al., 2022 and this study; combined). **c)** Dot plot showing all ‘molecular function’ GO-terms with adjusted p-values < 0.01 related to genes with upregulated mRNAs in *wago-1* gravid adult libraries. **d)** Dot plot showing all ‘molecular function’ GO-terms with adjusted p-values < 0.01 related to genes with downregulated mRNAs in *wago-1* gravid adult libraries.

## Footnotes

1 Reviewer access details: Log in to the PRIDE website using the following details: Project accession: PXD072057, Token: OxUIsCoIdzNV, Alternatively, reviewer can access the dataset by logging in to the PRIDE website using the following account details: Username, Password: diDHt44Y0B5U

